# Plasmid2MC: Efficient cell-free recombination of plasmids into high-purity minicircle DNA for use in genome editing applications

**DOI:** 10.1101/2024.07.08.602524

**Authors:** Roman Teo Oliynyk, Ahmed Mahas, Emil Karpinski, George M. Church

## Abstract

DNA plasmids are widely used for delivering proteins and RNA in genome editing. However, their bacterial components can lead to inactivation, cell toxicity, and reduced efficiency compared to minicircle DNA (mcDNA), which lacks such bacterial sequences. Existing commercial kits that recombine plasmids into mcDNA within proprietary bacterial strains are labor-intensive, yield inconsistent results, and often produce endotoxin-contaminated low-quality mcDNA. To address this challenge, we developed Plasmid2MC, a novel cell-free method utilizing **Φ**C31 integrase-mediated recombination to efficiently excise the bacterial backbone from conventionally prepared plasmids, followed by digestion of the bacterial backbone and all other DNA contaminants, resulting in highly pure and virtually endotoxin-free mcDNA. We demonstrated the application of mcDNA to express CRISPR-dCas9 for base editing in HEK293T cells and mouse embryonic stem cells, as well as for homology-independent targeted insertion (HITI) genome editing. The method’s ease of preparation, high efficiency, and the high purity of the resulting mcDNA make Plasmid2MC a valuable tool for applications requiring bacterial backbone-free circular DNA.

## Introduction

Plasmids, bacterial extrachromosomal self-replicating circular DNA elements, were first described by Joshua Lederberg in 1952 [1]. They were later found to facilitate sharing of antibiotic resistance and other inter-bacterial genome modification [2, 3]. Cohen et al. [4] demonstrated the construction of biologically functional bacterial plasmids in vitro in 1973. Since then, plasmids have been widely used for genome editing constructs delivery [5–7], including zinc-finger nucleases, transcription activator-like effector nucleases (TALENs), and CRISPR-based methods [8–12].

However, the prokaryotic sequences within plasmids, notably antibiotic resistance genes, are redundant for eukaryotic cell transfection and have the potential to elicit adverse effects on gene expression, cellular physiology, immune response [13–15], and lead to plasmid inactivation [15]. Minicircle DNA (mcDNA), produced from plasmids by eliminating bacterial backbones, can avoid these effects. mcDNA has shown lower inactivation, better nuclear uptake, and greater protein expression compared to plasmids [15–17], making it a promising tool for non-viral vector delivery [18].

Numerous studies have demonstrated the production of mcDNA using site-specific recombination [13, 14, 19, 20], where a serine integrase performs double-stranded DNA cleavage and subunit rotation to exchange DNA strands at two recombining sites present in a plasmid—the so-called attP and attB (abbreviated from attachment-phage and attachment-bacteria, respectively) sites [21]—thus separating the intended mcDNA and a bacterial backbone into two intercoiled DNA circles (see illustration in the Results). The recombination is then followed by a restriction enzyme digestion. Both recombinase and restriction enzyme genes are designed into specialized bacterial strains and an inducible activation is used to express them, resulting in the production of mcDNA within the bacteria [13, 14, 19, 20]. We found commercial kits [22] using this methodology labor-intensive and yielding inconsistent results. Purity and toxicity are known issues for such bacterially recombined mcDNAs, and additional cleanup is also cumbersome and labor intensive [18]. In a recent analysis, Almeida et al. [18] highlighted a common issue with specialized bacterial kits utilizing ΦC31 integrase and I-SceI: incomplete recombination and digestion, resulting in mcDNA tainted by remnants of parental plasmids and recombined empty bacterial backbone plasmids, which are unacceptable for therapeutic applications, posing significant challenges for downstream purification [23–25]. The FDA stipulates requirements for plasmids (and similarly mcDNA) intended for therapeutic use in vaccines, necessitating minimal bacterial genome and other DNA contamination, undetectable protein and RNA levels, and endotoxin levels ≤0.04 EU/*µ*g [26].

No protocol or commercial kit for cell-free plasmid recombination into mcDNA for use in genome editing experiments has been available. To address this shortcoming, we present Plasmid2MC, a cell-free protocol leveraging ΦC31 integrase—one of the best-characterized large serine recombinases to date [27–33]. In Plasmid2MC, ΦC31 recombination efficiently excises the bacterial backbone from conventionally prepared plasmids in a cell-free reaction, while spin column mcDNA extraction steps simultaneously reduce endotoxin levels, thus not requiring additional purification.

The ease of preparation, efficiency, and mcDNA purity of Plasmid2MC make it a superior alternative to existing kits on the market, such as those implementing inducible recombination and plasmid digestion within specially modified bacterial strains, produced by System Biosciences Inc. (SBI)—currently the only company offering such commercial kits. Our evaluation of SBI’s MC-Easy™ Minicircle DNA Production Kit revealed a costly, time- and labor-intensive process prone to multiple failure points and yielding mcDNA with high endotoxin levels and DNA contamination. Although recombination using ΦC31 integrase and other recombinases has been studied in laboratory experiments for decades, the Plasmid2MC method represents a novel, first-of-its-kind published protocol that delivers a simple and efficient cell-free approach for the practical production of high-quality mcDNA.

Finally, we demonstrated the efficacy of mcDNA prepared with Plasmid2MC method by using it to express CRISPR-dCas9 for base editing in HEK293T and mouse embryonic stem cells (mESCs), as well as for homology-independent targeted insertion (HITI) genome editing.

## Results

### Method overview

In this study, we developed an efficient cell-free method that takes as an input customarily cloned plasmids containing two recombining sites (attP and attB, Figure 1a), and with the help of ΦC31 integrase recombines these plasmids into two intercoiled DNA circles, one containing bacterial backbone with all components used for bacterial amplification, and another containing the mcDNA of interest Figure 1b,c.

**Fig. 1.**
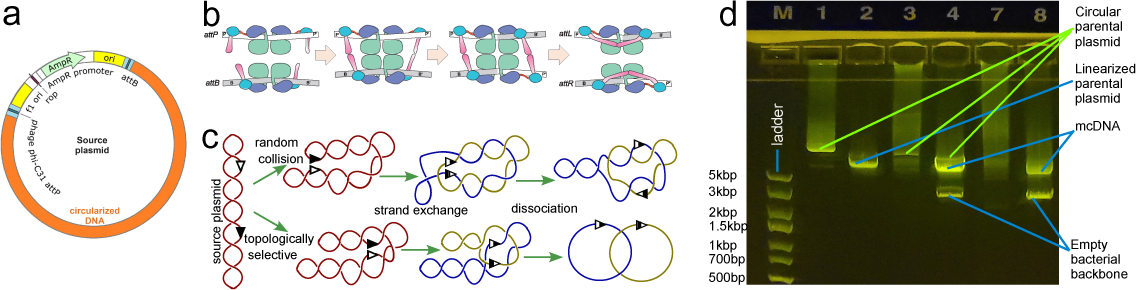
Φ**C31 recombination: mechanism diagram and experiment.** (**a**) Diagram of a plasmid containing attP and attB recombination sites separating the bacterial backbone and the intended mcDNA. (**b**) Subunit rotation mechanism for strand exchange by serine recombinases (this subplot is adopted from Rutherford et al. 2013 [34]). (**c**) Random collision and topologically selective strand exchange both lead to intercoiled structures of two DNA loops of varying complexity among the recombined molecules [21]. Random collision is the usual recombination mechanism, while topologically selective recombination was observed in experimental designs [21]. (**d**) Gel image showing the ΦC31 recombination of 10418 bp plasmid into a 2.7 kbp bacterial backbone circle and 7758 bp DNA circle. Each lane contains 300 ng of DNA. Lane M - DNA ladder; L1 - pure supercoiled plasmid; L2 - same plasmid with a single cut with a restriction enzyme (*Not*I); linear DNA travels faster than supercoiled DNA, and L1 presents a very weak matching band showing a minuscule amount of linear plasmid; L3 - after 80 minutes of recombination showing diffuse DNA scattering corresponding to the varying speed of travel of the miscellaneous catemers depicted in (B), and L4 same product cut with *Not*I, which is present in a single location on the bacterial backbone, thus allowing bacterial backbone release from the tangle, resulting in distinct bands (2.7 and 7.7 kbp) and a weaker band corresponding to the non-recombined plasmid; L7 and L8 - same as L3 and L4 after 12 hours of recombination; the original plasmid band is now nearly invisible (see the unprocessed gel photo in Supplementary Figure S1).

In the past, we developed a fully synthetic one-pot method for producing small circular DNA (up to 1000 bp) that uses only one spin column cleanup step (see our publication Oliynyk & Church [35]). We tried to apply the same one-cleanup approach to the Plasmid2MC method to avoid the inevitable loss of mcDNA during cleanup. However, preliminary experiments have shown that the cleanup step after the completion of recombination was indispensable in Plasmid2MC, even though it lowered the final mcDNA yield by estimated 16% for 5.7kbp source plasmid size and by 31.5% for 13.9 kbp source plasmid (see Methods DNA purification after the completion of recombination and Protocol steps). The recombination buffer containing 20 mM Tris, 0.1 mM EDTA, and 100 mM NaCl worked optimally (see Fine-tuning recombination buffer composition for optimal performance in the Methods), and experiments were performed with a recombination duration of 12 hours (see the finalized Protocol steps in the Methods section). The recombination reaction was then followed by a 1-hour Proteinase K digestion to remove the ΦC31 integrase, succeeded by the first QIAGEN QIAquick spin column cleanup of the recombined mcDNA, which was still intertwined with the bacterial backbone plasmid. QIAGEN advises using Proteinase K digestion for DNA samples with high protein content when using their DNA purificationspin column kits, which are highly selective against proteins [36]. This step indeed resulted in high-purity DNA with 260/280 ratios of 1.85 and 260/230 ratios of 2.17 for a 10.4 kbp source plasmid (see *Recombination Yield Table.xlsx* in Supplementary Data). The DNA recovery was on average 83.6% (a DNA loss of 16.4%) during the spin column cleanup for this plasmid size. Subsequently, we cleaved the empty bacterial backbone plasmid using the SspI restriction enzyme and degraded all linear and nicked DNA with T5 exonuclease, retaining only the intended mcDNA. This was followed by a second QIAquick spin column cleanup, yielding further purified mcDNA (see Supplementary Data and Methods section for detailed procedures).

### Plasmid2MC protocol yields at varying ΦC31 integrase and source plasmid DNA concentrations

The combined recombination results are presented in Figure 2 (see also Component concentrations in recombination experiments in the Methods section). Three plasmid sizes were used in the following analysis, titled as mcDNA size/source plasmid size in base pairs (bp)—3096/5758bp, 7758/10418bp, and 11282/13942bp—thus sampling the recombination protocol performance across a commonly used range of plasmid sizes and corresponding mcDNAs (see Plasmid design and assembly in the Methods, Supplementary Data, and Supplementary Figure S2). In Figure 2a,b, both ΦC31 integrase and source plasmid DNA (10418 bp plasmid used) had concentrations of 100 ng/*µ*l. Recombination efficiency gradually increases between 18 and 30°C, reaching 38% at 30°C, which is the optimal temperature (see Figure 2a). Recombination efficiency diminishes thereafter to 5% at 37°C. The reaction yield increased with longer recombination times, even showing additional gains from 33.3% at 4 hours to 39.4% at 8 hours of incubation, and remaining at a constant level thereafter (see Figure 2b).

**Fig. 2.**
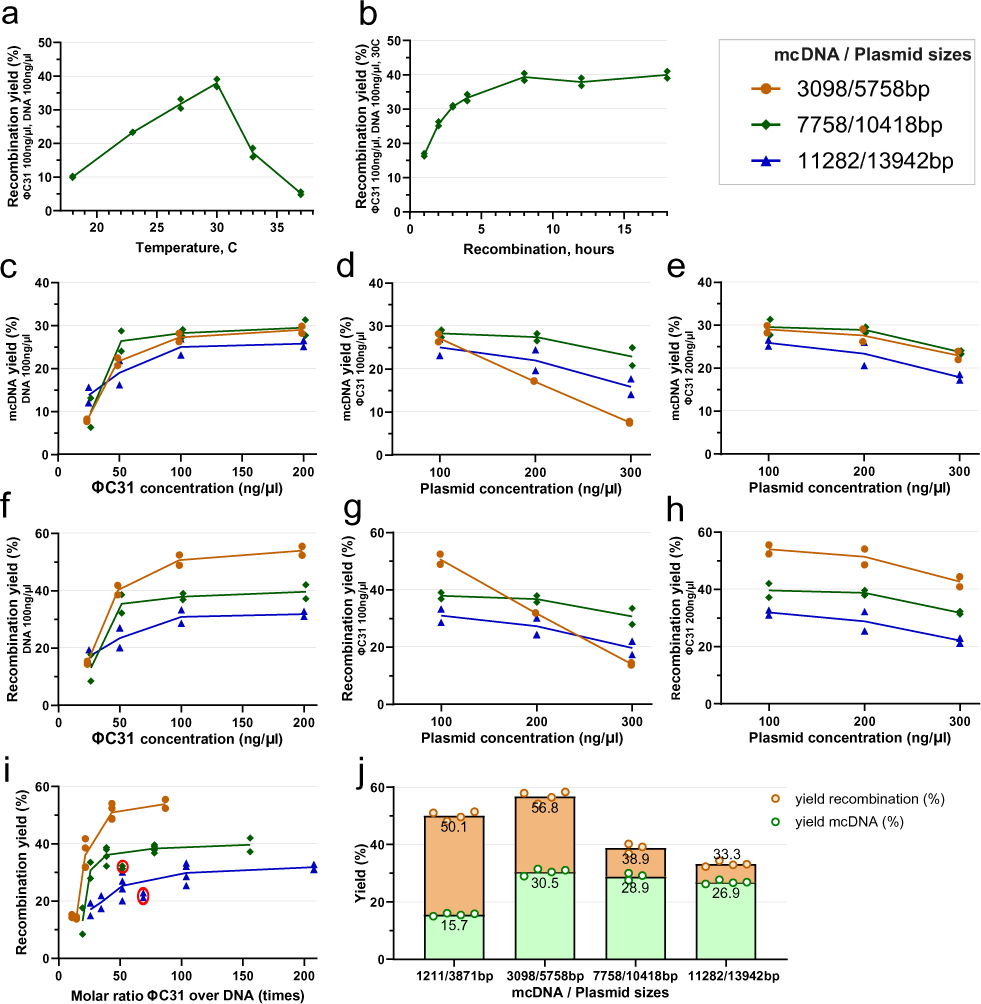
Recombination reaction efficiency and mcDNA yields. (**a**) Temperature curve of ΦC31 recombination efficiency, showing the distinct maximum yield at 30°C. (**b**) Recombination yield by time at the optimal temperature of 30°C. (**c**) mcDNA yield at ΦC31 integrase concentrations of 25–200 ng/*µ*l and a constant DNA concentration of 100 ng/*µ*l. (**d**) mcDNA yield at 100 ng/*µ*l ΦC31 and DNA concentrations of 100–300 ng/*µ*l. (**e**) mcDNA yield at 200 ng/*µ*l ΦC31 and DNA concentrations of 100–300 ng/*µ*l. The following plots (f)–(h) account for the bacterial backbone DNA circle for complete recombination efficiency (including length of bacterial backbone, and discounting processing losses): (**f**) Recombination yield at ΦC31 concentrations of 25–200 ng/*µ*l and a constant DNA concentration of 100 ng/*µ*l. (**g**) Recombination yield at 100 ng/*µ*l ΦC31 and DNA concentrations of 100–300 ng/*µ*l. (**h**) Recombination yield at 200 ng/*µ*l ΦC31 and DNA concentrations of 100–300 ng/*µ*l. (**i**) Combining the data points shown above presented a molar ratio of ΦC31 to source plasmid DNA molecules. (**J**) Maximum yield test at ΦC31 concentration of 200 ng/*µ*l and 20 *µ*g of input DNA at a concentration of 66.6 ng/*µ*l. The column titled 1211/3871bp represents yields for an additional 1211 bp mcDNA produced out of 3871 bp parental plasmid. Note: Tests for each data point were performed in duplicate for all subplots, with exception of (J) performed in quadruplicate and triplicate as seen in the subplot. Experiments in plots (B)-(J) were performed at 30°C and 12 hours incubation.

Further investigations with the three test plasmids at a constant plasmid DNA concentration of 100ng/*µ*l and variable ΦC31 integrase concentration (see Figure 2c) showed that at 25 ng/*µ*l of ΦC31, mcDNA yield was low for all three plasmids, gradually increasing with increasing concentrations, then beginning plateauing at 100 ng/*µ*l. An additional investigation with fixed ΦC31 integrase at 100 ng/*µ*l (see Figure 2d) showed a slight yield decrease for 10418 and 13942 bp plasmids when DNA concentration increased to 200 ng/*µ*l, with a steeper drop at 300 ng/*µ*l. At this ΦC31 integrase concentration, the yield for the shortest 5758 bp plasmid nearly halved from 27.2 to 17.2% when DNA concentration doubled; when DNA concentration tripled to 300 ng/*µ*l, the yield decreased to 7.6%. This suggests that a 100 ng/*µ*l ΦC31 integrase concentration may have been insufficient for the recombination of the short plasmid at higher DNA concentrations. Increasing the ΦC31 integrase concentration to 200 ng/*µ*l improved performance by 1–2% for all three source plasmids at a DNA concentration of 100 ng/*µ*l and mitigated the performance loss for the 5758 bp plasmid at higher DNA concentrations, resulting in parallel performance patterns for all three plasmids (see Figure 2e). This indicated that smaller plasmids, with a larger relative number of attP and attB sites per unit of weight, may require higher concentrations of ΦC31 integrase. Interestingly, the mcDNA yield was similar (25–30%) for the three plasmids at the most effective concentrations.

While the above results report the pure mcDNA yield, which are the values we measured directly, the successful recombination reaction includes a circular bacterial backbone of constant size, which was cut with a restriction enzyme and digested. Thus, the full recombination efficiency can be recalculated as:

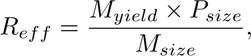

where *R_eff_* is the full recombination efficiency, *M_yield_* is the measured mcDNA yield, and *P_size_* and *M_size_* are the sizes of the plasmid and mcDNA, respectively. Figure 2f–H are equivalent to Figure 2c–e, showing the full recombination efficiency while accounting for the digested bacterial backbone according to the formula above. Recombination performances at concentrations of 200 ng/*µ*l of ΦC31 integrase and 100 ng/*µ*l DNA for the aforementioned experiments were 54%, 40%, and 32% for the shortest, medium, and longest plasmids, respectively. One explanation for this performance difference is that the cleanup column efficiency decreases with increasing DNA length, with column losses from two cleanups exceeding 50% for the largest source plasmid (see Protocol steps, Step 2). Additional DNA nicking or shearing may occur during initial plasmid extraction or intermediate cleanup, and these molecules will be digested by T5 exonuclease during the digestion stage. However, despite higher losses during cleanup for larger plasmids, the proportion occupied by the bacterial backbone in the larger plasmids was smaller than, for instance, in the 5758 bp plasmid, where the bacterial backbone made up nearly half of the plasmid size. This accounts for the similar mcDNA yield, despite the more significant differences in full recombination efficiency.

Recalculating the molar concentration of ΦC31 integrase and dividing it by the molar concentrations of the three test plasmids from the available data points (see Figure 2i, and Component concentrations in recombination experiments in the Methods) demonstrated that the recombination reaction yield is relatively low when ΦC31 molar excess is at or below 25X, and begins plateauing for all three plasmids at approximately 50X ΦC31 molar excess, with only fractional improvement at higher molar excesses. This figure is a sampling of clearly non-linear correspondence between concentrations and the molar excesses of these two variables. For example, at higher concentrations (DNA at 300 ng/*µ*l and ΦC31 integrase at 200 ng/*µ*l), the relative performance is notably below the trend line for lower molar ratios (see circled red data points for 10418 bp and 13942 bp plasmids in Figure 2i), showing that at such a high DNA concentration, the efficiency of recombination is diminishing.

The experiments described above were performed with elution volumes of 22–25 *µ*l, which aimed to achieve higher concentrations of DNA, thereby allowing for accurate measurements. Based on the observations in Figure 2i, we decided to verify the maximum yields at a concentration of ΦC31 integrase equal to 200 ng/*µ*l, with a reaction volume of 300 *µ*l and double the source plasmid weight of 20 *µ*g (DNA concentration of 66 ng/*µ*l), using an elution volume of 54 *µ*l. After incubating the columns at 55°C for 3 minutes before elution, centrifugation yielded 48–51 *µ*l of mcDNA eluate. The results presented in Figure 2j indicate that even with a higher elution volume, the recombination yield remained essentially identical to the results for ΦC31 integrase concentrations of 200 ng/*µ*l in Figure 2h. The 13942 bp source plasmid showed a 1.4% improvement, while the 10418 bp source plasmid exhibited a yield lower by 0.7%—within the inter-experimental variability. In this scenario, only the smallest 5758 bp source plasmid demonstrated a 3% improvement, resulting in a maximum recombination efficiency of 56.8% and an mcDNA yield of 30.5% at this plasmid size. This improvement was likely due to the molar ratio of ΦC31 integrase to DNA increasing from 87 in the first data point in Figure 2h to 130 for this source plasmid size.

This outcome aligns with our experience that heating the QIAquick cleanup columns at 55°C for 3–4 minutes before centrifugation results in an effective QIAquick column elution rate when using elution volumes of 20–25 *µ*l, with a repeated elution from the same column yielding only a few additional ng/*µ*l. This shows that the use of lower elution volumes, necessary for achieving high mcDNA concentration required for genome editing experiments, will not lead to significant mcDNA losses with QIAquick spin columns.

A question arose regarding the Plasmid2MC method’s capability to produce even smaller mcDNAs. To address this, we developed a shorter version of the plasmid, *p-att-SuperShort1211bp*, which measures 3871 bp and contains the same bacterial backbone, yielding mcDNA with a length of 1211 bp (see Methods). The yields of this construct are shown in the first column of Figure 2j, labeled 1211/3871bp (also see the sequencing diagram in Supplementary Figure S3). As anticipated, due to the smaller ratio of mcDNA size to the total plasmid size, the yield of mcDNA was relatively lower compared to longer constructs, at 15.7%. However, the overall recombination efficiency, accounting for all processing losses, remained high at 50.1%. The molar ratio for the reaction at the concentrations listed in Figure 2i was 88. It is expected that with an even smaller mcDNA-to-source-plasmid ratio, the mcDNA yield would decrease proportionally; nevertheless, smaller circular constructs can be produced more efficiently with our CV protocol, as further discussed in the Discussion section.

The high molar ratios of ΦC31 integrase raise a pertinent question: if only four molecules are sufficient for recombination per source plasmid, why do our experiments demand such an excess of integrase? Beyond the canonical ΦC31 attP and attB sequences (which do not exist verbatim in the human genome), approximately 27,924 pseudo-sites within the human genome can mimic these attP and attB functions [37]. These pseudo-sites facilitate the formation of functional ΦC31 dimers roughly every 107 kbp, a pattern potentially applicable to any DNA segment or plasmid. Moreover, the number of sites where a single ΦC31 molecule can bind, whether transiently or persistently, is significantly higher, contingent upon the binding energy of each site [38, 39]. Such off-target binding may limit the availability of exactly four ΦC31 molecules that need to simultaneously combine — two at attP and two at attB — for recombination to take place. In Supplementary Figure S21, we reanalyzed data from Figure 2I to estimate the number of ΦC31 molecules per kbp of source plasmid. The occurrence of these single-integrase binding sites varies with plasmid DNA sequence, and indeed Supplementary Figure S21 indicates that recombination efficiency plateaus when exceeding five ΦC31 molecules per kbp, with all tested plasmids showing diminished recombination below 2.5 ΦC31 molecules per kbp.

### Optimal concentrations of source DNA and ΦC31 integrase

Having consolidated the above data, we can formulate the consideration for the optimal concentrations and quantities of ΦC31 integrase and source plasmids. Plasmid Midipreps typically yield 400–600 *µ*g of plasmid per 45ml of *E. coli* culture, providing abundant source plasmid DNA. To economize the use of ΦC31 integrase, it may be optimal to use 100 ng/ul of ΦC31 and 200–300 ng/*µ*l of DNA for larger plasmids (e.g., see Figure 2d,g) and a somewhat higher ΦC31 concentration for smaller plasmids (see the 5758 bp plasmid in Figure 2e,h).

While it is practical to measure concentrations in ng/*µ*l, it is important to keep in mind that the molar ratio (or recalculate in less common units of ΦC31 molecules per kbp of source plasmid) is a crucial factor for determining the quantities used in Plasmid2MC reactions.

### Collateral reduction of endotoxin levels in mcDNA compared to source plasmids

Although no specialized cleanup steps were employed beyond the two spin column purifications required for Plasmid2MC processing, we observed a significant reduction in endotoxin levels. This finding warranted a more detailed investigation, described below. Tests using the Endosafe® nexgen-MCS endotoxin testing equipment revealed that source plasmids prepared with the QIAGEN Midiprep Plus kit had endotoxin levels ranging from 0.17 to 1.46 EU/*µ*g of DNA. While the QIAGEN kit specifies endotoxin levels below 1 EU/*µ*g, one of three samples slightly exceeded this limit (see endotoxin testing reports in Supplementary Figure S20 and preliminary data in Supplementary Figure S15). To assess a scenario with slightly elevated initial endotoxins, we selected a parental plasmid with a level of 1.46 EU/*µ*g of DNA and determined that the Plasmid2MC protocol lowered the endotoxin content in the resulting mcDNA to 0.042 EU/*µ*g—a 35-fold reduction (see Figure 3A). Notably, although the protocol incorporates two spin column cleanups with washing steps, these alone do not suffice for substantial endotoxin removal. We hypothesized that endotoxin reduction occurs during the digestion stage, where T5 exonuclease and the *Ssp*I restriction enzyme bind endotoxins, forming protein-endotoxin complexes [40, 41] that are subsequently removed during washing. To test this, we used the plasmid p-SBkit-ABE8e-parental-plasmid.gbk, which lacks a recognition site for the *Ssp*I restriction enzyme (see Figure 3b and corresponding endotoxin testing reports in Supplementary Figure S17). Incubating 10 *µ*g of this plasmid per step 3 of the Plasmid2MC protocol reduced endotoxin levels 4-fold, from 0.170 EU/*µ*g to 0.039 EU/*µ*g after cleanup. However, introducing Proteinase K after the digestion step resulted in no improvement of endotoxin levels, which remained close to the initial level. Thus, using Proteinase K after the recombination reaction is crucial (see Materials and methods), but proteinase digestion should not follow T5 exonuclease and restriction enzyme digestion.

**Fig. 3.**
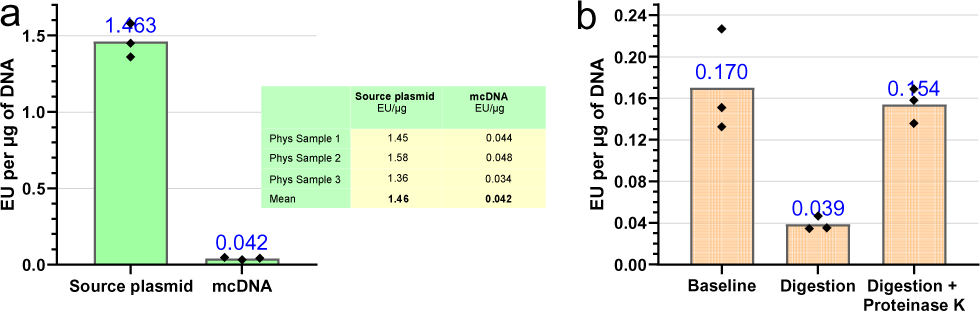
Reduction of Endotoxin Levels During Plasmid2MC Processing. **(a)** A 35-fold reduction in endotoxin concentration was observed from the source plasmid to the resultant mcDNA (see endotoxin testing reports in Supplementary Figure S16). **(b)** Digestion experiments supported the hypothesis that proteins involved in digestion might bind endotoxins, facilitating their removal during the Plasmid2MC spin column purification step (see endotoxin testing reports in Supplementary Figure S17).

### Comparative evaluation of the System Biosciences Inc. MC-Easy™ Minicircle DNA Production Kit

A detailed comparative analysis of the commercially available SBI MC-Easy™ kit is presented in the Supplementary Chapter titled ‘Comparative evaluation of System Biosciences Inc MC-Easy™ Minicircle DNA Production Kit,’ with key results summarized in Supplementary Figures S11–S14,S18. Testing showed that the recommended procedure produced mcDNA with endotoxin levels ranging from 47 to 121 EU/*µ*g, accompanied by detectable traces of parental plasmid, recombined empty bacterial backbone, and *E. coli* genomic DNA (see Supplementary Figures S11–S12). While the optional DNase digestion and subsequent cleanup steps recommended by SBI effectively eliminated DNA contamination, they also led to a significant loss of mcDNA. Even after this additional purification, endotoxin levels in mcDNA generated by the SBI MC-Easy™ Kit were reduced by only approximately 3-fold, remaining high at 14.2–45.8 EU/*µ*g. Beyond these elevated endotoxin levels, the SBI MC-Easy™ kit is expensive (see pricing comparison in the Supplementary and in Discussion section) and demands a 3–4-day preparation and quality analysis process, featuring at least four major failure points—if any fail, the entire minicircle preparation must be restarted. Although the patented SBI MC-Easy™ approach may appear clever at first glance (see Supplementary Information for details), its limitations likely arise from recombining and digesting plasmids within living *E. coli* cultures—a method proven too complex for consistent efficacy and prone to significant endotoxin contamination due to the large volumes (400 mL) of bacterial culture required, which ultimately yield relatively low amounts of mcDNA.

**In Comparison**, the Plasmid2MC method employs routine bacterial amplification of the source plasmid. Subsequent steps, outlined in the Methods section, are conducted entirely in a cell-free, in vitro system, enabling precise timing in a thermocycler and eliminating the failure points inherent in the less reproducible, monitoring-intensive bacterial production process of the SBI MC-Easy™ Kit. As a result, Plasmid2MC is faster, more cost-effective, and more reliable, producing notably higher-quality mcDNA than the MC-Easy™ Kit. With endotoxin levels as low as 0.042 EU/*µ*g, Plasmid2MC-derived mcDNA is exceptionally well-suited for transfection into sensitive cell lines.

### The mcDNA use in CRISPR-dCas9 base editing in HEK293T and mESCs

Having optimized the protocol using ΦC31 integrase for the effective production of mcDNA, it is customary to demonstrate the performance of the resulting DNA constructs. For this validation, we used the plasmid *p-att-ef1a-ABE8e-dCas9-BSD* (see the Materials and methods) and corresponding mcDNA produced using our protocol. This plasmid is an implementation of TadA for the base editing version ABE8e introduced by Richter et al. [11], assembled into our plasmid containing ef1*α* promoter, attP and attB sites, blasticidin selection resistance gene, and dCas9—which is used in our ongoing research (rather than nCas9 used by Richter et al. [11]; see Figure 4a). Applying the Plasmid2MC protocol to the source plasmid *p-att-ef1a-ABE8e-dCas9-BSD*, we produced mcDNA for use in these validation experiments. The resulting mcDNA was primarily represented by a single circular DNA, confirming the known ΦC31 integrase preference for intra-molecular recombination [21], with a small proportion of double-sized intermolecular concatemers, marked “dimer” on the sequencing histogram from Plasmidsaurus (see Figure 4b, Supplementary Figure S2). The sequencing showed the perfect matching of the expected and achieved mcDNA (sequencing files are available in the Supplementary Data).

**Fig. 4.**
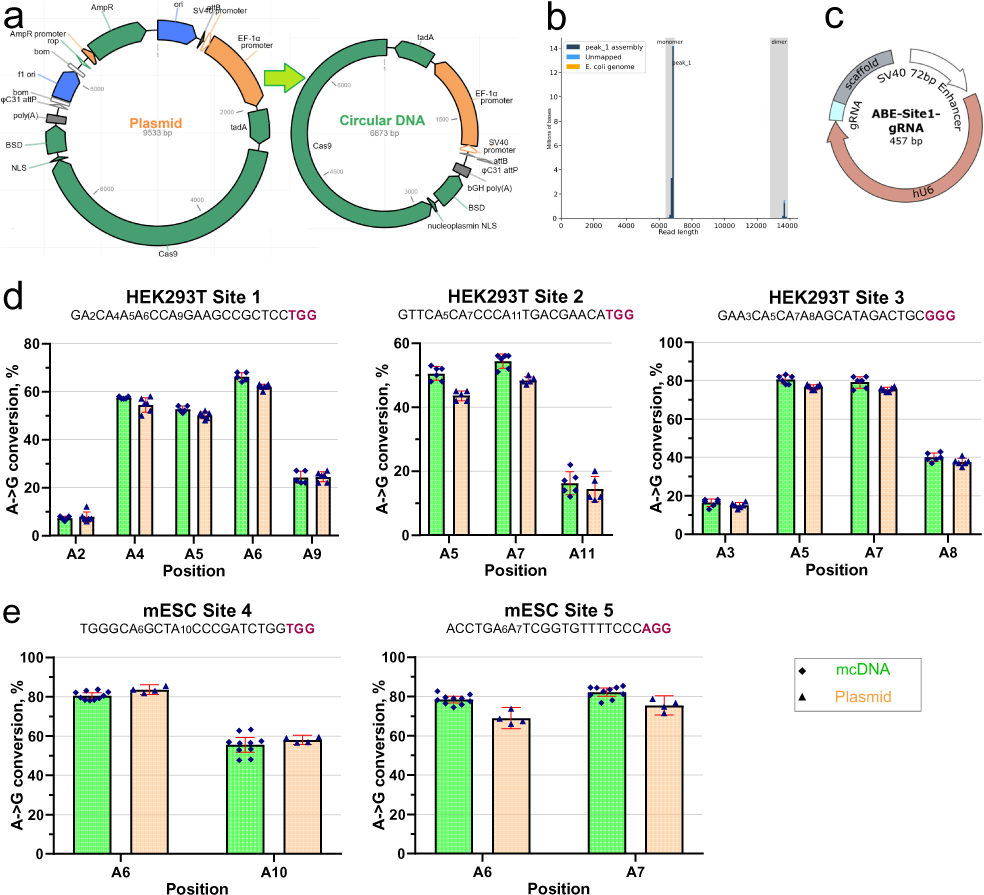
Validation of mcDNA performance in ABE8e dCas9 base editing. (**a**) Diagram of the source plasmid and the resulting mcDNA with excluded bacterial backbone, used in this validation edit. (**b**) Plasmidsaurus sequencing histogram showing the high quality of the mcDNA product. (**c**) A circular vector for the expression of gRNA for base editing targeting, using the method developed earlier in our laboratory [35]. (**d**) Base editing (dCas9 ABE8e) A*→*G conversion rates in HEK293T cells for the first three sites from [11]. (**e**) dCas9 ABE8e editing A*→*G conversion rates in two mESC sites. Colors: mcDNA (green) and the original plasmid (orange). Guide sequences for the sites are followed by PAM (in bold red font).

We used the Circular Vector (CV) protocol to prepare the five gRNA expression vectors used for five base editing experiments (see Supplementary Figure S4). Briefly, the CV protocol takes a linear double-stranded DNA fragment and ligates/purifies it into short, circularized DNA containing the gRNA expression element, as illustrated in Figure 4c, which can be completed and ready for transfection in as little as 3 hours (for implementation details, see Oliynyk & Church [35]). This protocol is particularly useful for making multiple gRNA expression vectors with minimal hands-on time, especially when used in conjunction with larger plasmids expressing proteins (and now mcDNAs), which are more labor-intensive but can be reused for multiple editing targets. We targeted three sites from Richter et al. [11], demonstrating A→G conversion in HEK293T cells (see Figure 4d). Figure 4e also demonstrates successful A→G conversion at two locations in mESCs using the same plasmid and mcDNA. The editing of both HEK293T and mESCs was followed by antibiotic (blasticidin) selection. The editing performance was similar in magnitude and differed only slightly between the plasmid and the mcDNA. The editing efficiency for HEK293T cells was consistently high, due to our extensive experience with this cell line, and additional attempts at optimizing transfection and selection parameters resulted in minimal improvement. However, after several trials, we succeeded in optimizing the transfection and selection of mESCs (see Methods), achieving an editing efficiency above 80%. We performed NGS sequencing to validate these results, and indeed, the editing efficiency was even slightly higher than what was observed in EditR Sanger sequencing analysis in Figure 4e, showing 86.5% editing for position A6 and 81.7% editing for position A10 for mESC Site 4 (see Supplementary Figure S7). These experiments showed comparable base editing performance when using the plasmid and mcDNA in both cell types, as should have been anticipated in such basic validation experiments. Maniar et al. [16] demonstrated minicircle inactivation in non-dividing liver cells starting four days after transfection, with increasing inactivation thereafter. By this time, the dilution in concentrations of both mcDNA and plasmids in HEK293T cells, and even more so in mESCs, would far outweigh the eventual inactivation of expression vectors in terms of base editing efficiency.

### The mcDNA use in homology-independent targeted integration (HITI)

mcDNAs have garnered significant interest due to their utility in site-specific genome integration applications. HITI utilizes mcDNA for the insertion of exogenous DNA into targeted genomic sites without the need for homologous recombination [42, 43]. Here, we further demonstrated the utility of the Plasmid2MC method to produce HITI-compatible mcDNAs. We designed mcDNA constructs encoding a fluorescent protein cassette (mKate) linked to an antibiotic selection marker (Hygromycin) via a T2A peptide sequence, which allows for the in-frame insertion of both the fluorescent protein and the selection marker into housekeeping genes, including ACTB and GAPDH (see Figure 5a). To showcase the preparation of larger HITI templates, we added a non-functional random DNA sequence downstream of the hygromycin resistance gene, increasing the total size of the HITI template to approximately 3 kbp. The integration sites were strategically chosen just upstream of the stop codons of these housekeeping genes. A P2A sequence was incorporated immediately upstream of the donor template sequence to ensure that the insertion of the HITI constructs occurred in-frame, facilitating the translation of the downstream HITI elements as part of the same polypeptide as the endogenous proteins (see Figure 5a). We employed the CRISPR-Cas9 system with gRNA CVs specifically designed to target these integration sites alongside our mcDNA constructs. Following transfection, cells were cultured under hygromycin selection for two weeks to ensure only those cells that successfully integrated the mcDNA constructs survived (see Figure 5b).

**Fig. 5.**
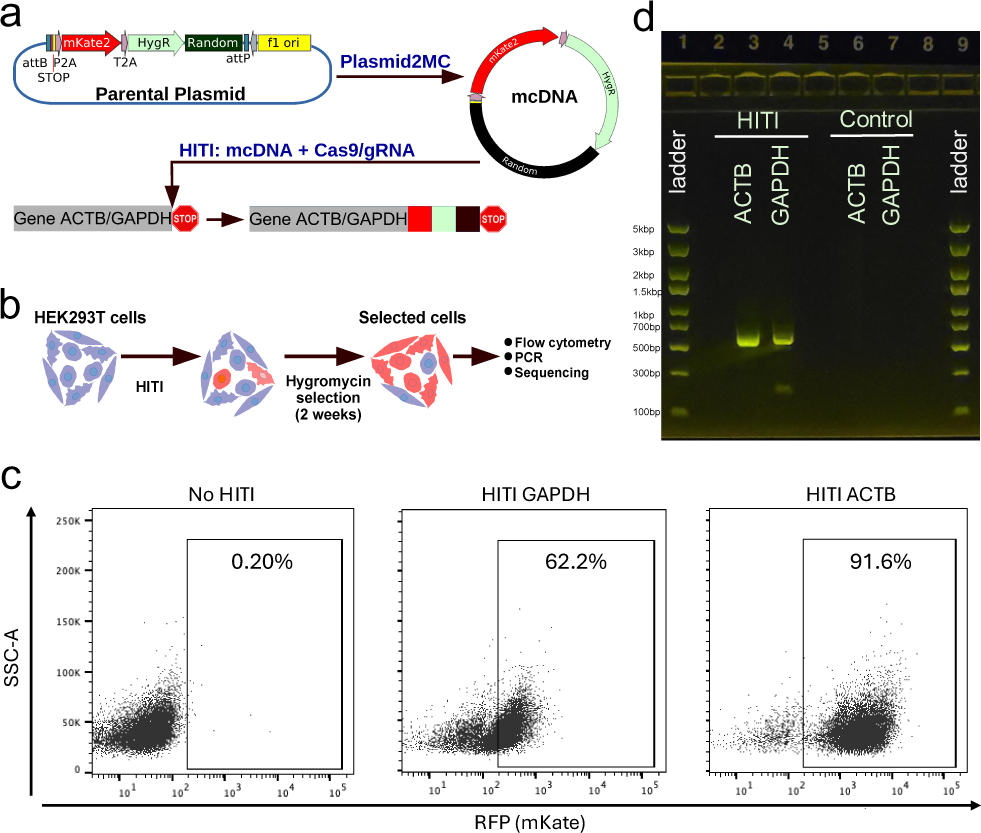
HITI editing results. (**a**) Schematic showing the design of HITI constructs. Blue pentagon, Cas9/gRNA target sequence in the host genome and HITI mcDNA. HygR: Hygromycin-resistant gene. Black square: non-functional random DNA sequence. (**b**) Schematic of HITI experiment with antibiotic selection to enrich cells with the correct HITI-mediated integration before subsequent analysis. (**c**) Flow cytometry analysis of cells treated with HITI or control (untransfected cells). Percentage of positive population is shown within the gates. (**d**) Gel electrophoresis of PCR overlapping the mKate gene and either the ACTB or GAPDH genes. HITI - shows bands for edited samples, while no bands are visible for unedited Controls. See Supplementary Figure S9 for additional analysis and unprocessed gel electrophoresis photos.

Flow cytometry analysis revealed a significant population of cells expressing mKate, with 91.6% of ACTB-targeted cells and 62.2% of GAPDH-targeted cells displaying the red fluorescent protein, indicating successful integration and expression (see Figure 5c). To confirm the integration of the HITI mcDNA at the genomic level, PCR was conducted using primers specific to the mKate gene and either the ACTB or GAPDH genes. The expected amplicons were observed only in HITI-treated cells, with no amplification in control groups (see Figure 5d). Subsequent Sanger sequencing of these amplicons verified the precise integration of the HITI constructs at the targeted genomic loci, confirming the correct junctions between the mcDNA and the host genome. These findings substantiate the efficacy of the Plasmid2MC method for producing large HITI-compatible mcDNA donor templates.

## Discussion

The Plasmid2MC method and protocol presented here elucidate the preparation of mcDNA from conventionally amplified or cloned plasmids containing ΦC31 attP and attB sites at the junctions between the bacterial backbone and the intended mcDNA, facilitated by ΦC31 integrase. In this protocol, ΦC31-mediated recombination efficiently excises the bacterial backbone, followed by purification steps that digest the separated bacterial backbone and reduce endotoxin levels (see Materials and methods). Our approach provides an alternative to costly proprietary kits that rely on specialized bacterial strains, yielding highly purified mcDNA with low levels of bacterial endotoxins.

Notably, the intrinsic recombination reaction efficiency approaches 100%, with DNA losses primarily occurring during these purification steps, resulting in an overall protocol efficiency ranging from 56% to 34% for plasmid sizes ranging from 5.8 to 13.9 kbp, respectively. Using a 2.7 kbp bacterial backbone, the pure mcDNA conversion efficiency is an impressive 27% to 30% after accounting for the backbone size. Applying the Plasmid2MC method to produce a short 1211 bp mcDNA revealed that the recombination efficiency was maintained at 50.1%. However, due to the smaller ratio of mcDNA to the entire parental plasmid, the yield of mcDNA was reduced to 15.7%, which might still be practical in certain scenarios. For even shorter circular DNA constructs, the CV method, discussed further below, could prove more efficient. Although a conversion yield of 34–56% might initially seem low, it’s important to consider that we use conventionally produced, inexpensive, and plentiful plasmids as starting material. The advantage here is the straightforward and efficient production of high-purity mcDNA in quantities suitable for genome editing experiments.

Our current pricing for a single custom project, which yielded 18.96 mg of ΦC31 integrase, was quoted at $1,552.95 by GenScript. At such a price, the custom order is less expensive than a 10-prep MC-Easy™ kit, which costs $1,914 (Cat# MN925A-1) from System Biosciences Inc. With a recombination yield of 30%, this quantity of ΦC31 integrase would be sufficient to produce at least 5.6 mg of mcDNA, and potentially up to double that amount by using a higher excess of source plasmid to conserve ΦC31 integrase, making ordering a large quantity of*P hi*C31 cost-effective. Currently, ΦC31 protein is not available off-the-shelf in small quantities; however, one can anticipate its availability materializing with demand, akin to many commercially available proteins and restriction enzymes. Hypothetically, a kit containing 1% of this amount of ΦC31 integrase would cost $15.53 and could produce between 56–112 *µ*g of mcDNA. Assuming that the production of ΦC31 integrase at industrial volumes would cost significantly less than what we paid, and considering that the recombination buffer contains no costly or hard-to-handle components, such a kit could be priced only slightly above this estimate. Even if the price of this hypothetical ΦC31 integrase kit were to quadruple from this estimate, it would still be competitively priced compared to many commercially available restriction enzyme and protein kits. Until such time, contracting a larger quantity of ΦC31 from a manufacturer like GenScript is an easy and cost-effective option, with the only drawback a longer delivery time.

There are fundamental differences between the Plasmid2MC method and our previously published CV method [35], which is primarily aimed at efficient fully synthetic substitution for plasmid-based guide RNA expression in CRISPR, base, and prime gene editing in cell cultures. The CV method offers rapid cell-free preparation of short circular DNA ready for transfection within 3 hours after receiving dsDNA from a commercial supplier, with minimal hands-on time [35]. In Oliynyk & Church 2022 [44], we noted that error rates in commercially delivered dsDNA fragments were ∼1/6,000 per nucleotide for various commercial products [45–47]. This error rate is excellent for short CRISPR-Cas9 guide RNA sequences but may lead to a significant proportion of defective proteins when extending the circular DNA length to thousands of base pairs [44] in the case of mcDNA. Additionally, the CV method performance is optimal at a CV size of 450 bp, where it reaches up to 62% efficiency in converting input linear double-stranded DNA into the circular expression vectors, diminishing to 30% efficiency at a 950 bp CV size and further to 5–8% efficiency at 1800 bp. This led us to define 450–950 bp as the practical range of the CV protocol [44]. The principal reason for this is that at longer linear source DNA sizes at practical concentrations, the CV ligation reaction favors polymerization into infinite chains of concatemers, with a vanishingly low proportion of circular DNA units [48, 49].

The Plasmid2MC method addresses the aforementioned limitations of the CV method by leveraging *E. coli* amplification of a single plasmid clone, virtually eliminating DNA replication errors [44, 50, 51]. Our sequencing of recombined and purified mcDNA invariably showed a perfect match of the expected and achieved mcDNA sequences. Consequently, mcDNA produced by this method proves applicable not only for protein-expressing genome editing constructs but also for HITI templates and other large circular DNA constructs. Both methods can be used complementarily, as demonstrated in this study where CVs are utilized for guide gRNA expression alongside Cas9 and other proteins expressed by mcDNA in base editing validations. The Plasmid2MC method enables the production of larger quantities of mcDNA, which can be reused across multiple experiments, while the CV protocol efficiently generates multiple guide RNAs as needed. Our lab continues to explore genome editing by combining these two approaches.

In most scenarios, kits like the QIAGEN Midiprep Plus ensure the removal of proteins and RNA during source plasmid preparation, whereas the Plasmid2MC protocol not only lowers endotoxin levels but also fully digests all unintended DNA. The FDA sets stringent requirements for plasmids and mcDNA intended for therapeutic applications in vaccines, demanding minimal bacterial genome and other DNA contamination, undetectable levels of protein and RNA, and endotoxin levels ≤0.04 EU/*µ*g [26]. While FDA-certified mcDNA production would require well-documented equipment, procedures, and testing of each production lot by a medical or pharmaceutical provider, which was outside the scope of our research, the Plasmid2MC method can be used as a starting point for establishing such production. The Plasmid2MC method is capable of producing mcDNA that closely approaches these requirements, evidenced by a 35-fold reduction in bacterial endotoxins when using conventional plasmids produced with the help of QIAGEN Midiprep Plus as starting material. In contrast, we found orders of magnitude higher endotoxin levels in the mcDNA produced using the System Biosciences Inc MC-Easy™ Minicircle DNA Production Kit.

In conclusion, the method’s ease of preparation, efficiency, and mcDNA purity make Plasmid2MC a valuable new tool for CRISPR-Cas9 and other applications that require bacterial backbone-free circular DNA.

## Materials and methods

### Plasmid2MC protocol

The flow of the Plasmid2MC protocol is presented in Figure 6.

**Fig. 6.**
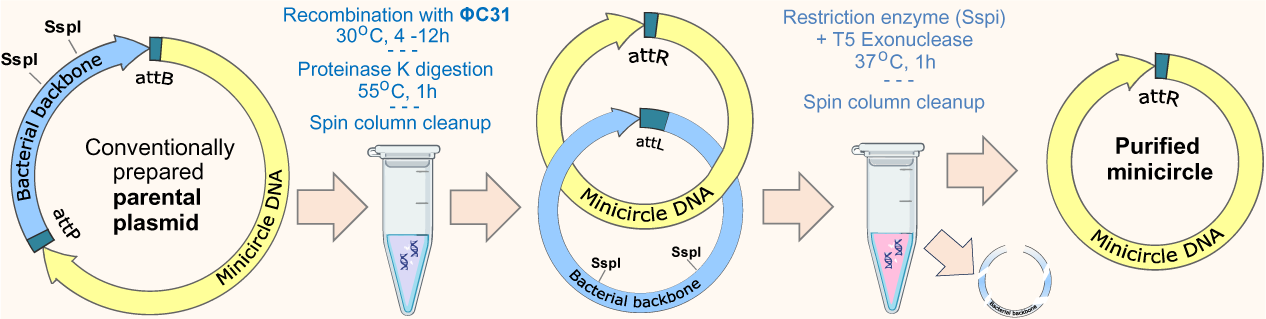
Plasmid2MC protocol flow.

#### Preliminary steps

Most of the reaction enzymes and buffers were based on commercially available kits and commonly used equipment. The exceptions include obtaining ΦC31 integrase and preparing the 2X reaction buffer as described below, and a small modification to the design of the plasmids.

1. **Obtain** Φ**C31 integrase protein from a reliable vendor**. We subcontracted GenScript and received 18.96mg of ΦC31 protein at a concentration of 4.74 mg/ml (see Reagents and Materials).
2. **Dilute the** Φ**C31 integrase with pure glycerol to make a working buffer**. Add glycerol to achieve a 50% vol/vol concentration suitable for storage at -20°C in liquid form, sufficient for a few months of experiments and store the remaining ΦC31 integrase at -80°C for long term storage. In our case, the ΦC31 integrase concentration in the working buffer became 2.6 *µ*g/*µ*l.
3. **Prepare the 2X recombination buffer to contain:** 200 mM NaCl, 40 mM Tris pH, 0.2 mM EDTA pH 7.5.

For example, 50ml of 2X buffer can be prepared as follows:

a) 10 ml of 10mM-Tris, 1mM-EDTA (pH 8.0; Fisher Scientific Cat#BP2473) and a negligible addition of Tris to produce 2X of 0.1 mM EDTA.
b) 2 ml of 1 M Tris (pH 8.0; Corning Cat#46-031-CM).
c) 584 mg of NaCl.
d) 38 ml of nuclease-free *H*_2_*O* (Invitrogen 10977015).

Note: The literature lists a variety of buffer compositions and additives, such as TPP, BSA, and spermidine [21, 27, 52]. Our testing demonstrated that customarily used ingredients other than those listed above resulted in no effect—or even lower recombination yield—and appeared to be unnecessary. As demonstrated in the Results, we achieved complete recombination using our 2X buffer in overnight reactions, while ΦC31 protein was sufficiently stable to provide yield improvements over a 8-hour reaction versus a 4-hour reaction. Buffers with NaCl concentration in the range of 100–150 mM worked equally well, and the small increase of NaCl from the working buffer remained well within this range, even at the highest concentration of ΦC31 integrase (the Φ31 integrase storage buffer delivered by GenScript contained a relatively high NaCl concentration, see Reagents and materials).

4. Make alterations to the design of source plasmids to include ΦC31 attP and attB sequences and add or identify unique restriction enzyme sites in the bacterial backbone area. The sequences are attB 5’-GTGCCAGGGCGTGCCCTTGGGCTCCCCGGGCGCG and attP 5’-GCCCCAACTGGGGTAACCTTTGAGTTCTCTCAGTTGGGGG, with the desired for circularization DNA sequence in between. Position them in a manner that allows the separation of the desired mcDNA and bacterial backbone elements into two circular units. The recombination occurs near the middle of the above sequences, with circularized DNA containing 5’-GCCCCAACTGGGGTAACCTTTGGGCTCCCCGGGCGCG, which may need to be considered when designing the mcDNA for a specific purpose.

The bacterial backbone must contain a unique restriction enzyme site, which will be used to cut and digest the bacterial backbone while leaving the desired mcDNA intact. It is preferred to use an efficient restriction enzyme. As only three examples out of many possibilities, we prefer to use New England Biolabs (NEB) SspI-HF, NotI-HF, or BsaI-HFv2 since these are highly efficient and inexpensive enzymes. In this study, the bacterial backbone contained two *Ssp*I restriction sites, and we report the results using SspI-HF.

Clone the plasmids as usual [53–55], amplify using a competent *E. coli*, and extract DNA using a suitable kit for the required quantity of plasmid DNA. You can use our plasmid (deposited at Addgene as p-att-ef1a-ABE8e-dCas9-BSD #226116) for a bacterial backbone containing convenient restriction enzyme sites around attP and attB, as well as double *Ssp*I restriction enzyme sites and a unique *Not*I restriction enzyme site within the backbone. The sequence is also available in the Supplementary Data (*p-att-ef1a-ABE8e-dCas9-BSD.gbk*).

#### Reagents and materials

A detailed inventory of the reagents and resources used in this research can be found in Supplementary Table S1.

ΦC31 integrase protein was subcontracted from GenScript. We received 18.96 mg of protein, the currently quoted pricing for such order is $1,660.00. The product concentration was 4.74 mg/ml(by Bradford) A260/A280: 0.557, ≥85% purity, endotoxin level 0.338 EU/mg or equivalently 0.000338 EU/*µ*g effectively endotoxin free. See the protein amino acid sequence and QA in Supplementary Figure S10.

The GenScript ΦC31 integrase storage buffer composition was: 50 mM Tris-HCl, 500 mM NaCl, 10% Glycerol, pH 8.0.

#### Protocol steps

1. **Recombine the source plasmid.** Prepare the recombination reaction as listed in Table 1 below, with ΦC31 integrase and 2X buffer being half the reaction volume and plasmid DNA in water the other half. Recombine for at least 4 hours, preferably 8–12 hours or overnight. It is necessary to add proteinase K and digest ΦC31 integrase after the completion of recombination because both intact and degraded ΦC31 integrase negatively affect the column’s efficiency (consult the further section DNA purification after the completion of recombination).
2. **Use the spin column kit to clean up the recombined DNA.** Use a spin column purification kit to perform intermediate purification of the recombination reaction output. We used QIAGEN’s *QIAquick PCR Purification Kit* [56]. Although the kit advertises use with up to 10 kbp DNA size, we found that it still performed well and resulted in cleaner DNA at 13.9 kbp when compared to other kits. Ensure you perform Proteinase K digestion on the ΦC31 integrase to prevent interference with DNA binding to the purification columns. For improved elution yield, heat the spin column according to the manufacturer’s recommendations. We warmed the purification columns in a heat block at 55°C for 3 minutes before elution. For maximum elution efficiency, we used ≥60 *µ*l of water. It is not necessary to measure the concentration of the eluted DNA at this step, unless desired. Typical elution yield is approximately 90% for short DNA fragments, but significantly less for larger plasmid sizes (see the note below). Use multiple columns if needed, based on the column binding capacity.

Note: We validated the efficiency of the purification kits used for a similar purpose in Oliynyk & Church [35, 44], where we preferred QIAGEN’s *QIAquick PCR Purification Kit* (QIAquick) [56]. In this research, we additionally tested the yield of QIAquick and Takara’s *NucleoSpin Gel and PCR Cleanup* (NucleoSpin) [57] with a 13.9 kbp plasmid. We found that with the input of 10 *µ*g of the largest (13.9 kbp) plasmid used in this research, we retrieved 7.46 *µ*g (74.6%) with NucleoSpin and 6.85 *µ*g (68.5%) with QIAquick, using an elution volume of 55 *µ*l in both cases. However, we found that when we used both kits for two-stage purification, the yield of NucleoSpin was lower, while the NucleoSpin DNA quality with preheating recommended by the vendor was consistently showing salt carry-over, with 260/230 quality typically in the 1.5–1.8 range, while the production runs of QIAquick exhibited excellent DNA quality. We achieved an even higher efficiency of *>*85% large DNA retrieval rate using the QIAGEN QIAEX II Gel Extraction Kit; however, the DNA produced by this kit appeared to carry over resin particle pollution, and contamination indicated by an absorption peak at 225 nm with an amplitude exceeding the 260 nm DNA peak. QIAGEN does not recommend using DNA produced by QIAEX II kit for transfection, but recommends using the QIAquick kit instead, and we concur. Thus, we opted to use a QIAquick kit for a better combination of mcDNA final yield and quality. The two cleanups required have an estimated combined retrieval of the plasmid and mcDNA of 0.685 x 0.685 = 0.469 (46.9%)—i.e., 53.1% is lost in the column cleanup for this DNA size. This implies that with the maximum reported reaction yield of 33.9% for 13942 bp source plasmids—and assuming that recombination reaction efficiency is close to 100%—the additional pipetting losses and digestion of nicked and broken DNA constitute the remaining 13.0% difference.

**Table 1.**
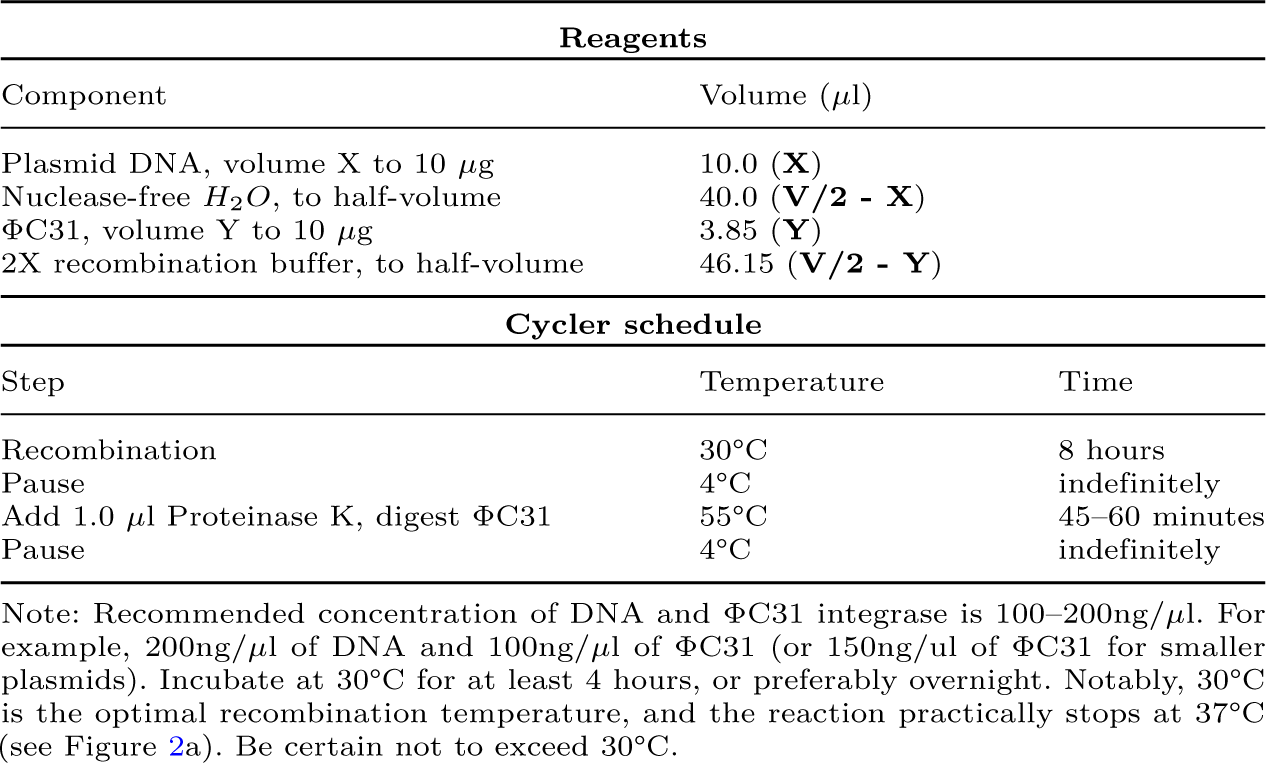
Recombination reaction example for 100 ng/*µ*g (10.0 *µ*g) of input of both plasmid DNA at 1.0 *µ*g/*µ*l and ΦC31 at 2.6 *µ*g/*µ*l in volume V = 100 *µ*l.

3. **Cut the bacterial backbone DNA and digest it and all other non-circular or nicked DNA.** Perform the reaction (as listed in Table 2) using a restriction enzyme unique to the bacterial backbone.

**Table 2.**
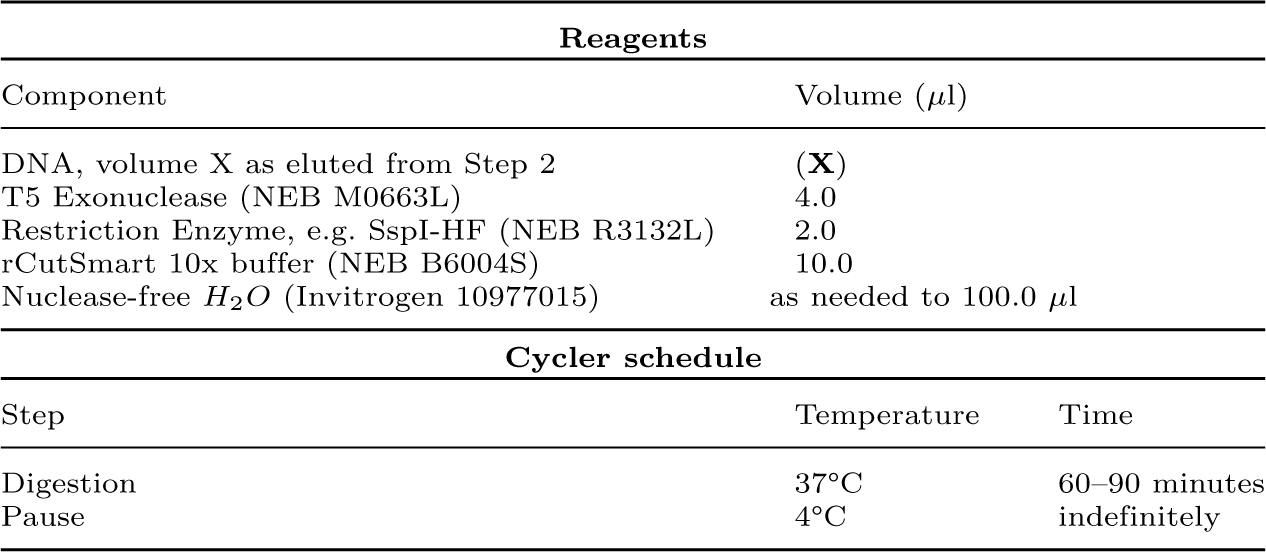
Example of the digestion/purification recombined DNA; reaction volume V = 100 *µ*l.

| Reagents |  |  |
| --- | --- | --- |
| Component | Volume ($\mu\text{l}$) | |
| DNA, volume X as eluted from Step 2 | (X) |  |
| T5 Exonuclease (NEB M0663L) | 4.0 |  |
| Restriction Enzyme, e.g. SspI-HF (NEB R3132L) | 2.0 |  |
| rCutSmart 10x buffer (NEB B6004S) | 10.0 |  |
| Nuclease-free $\text{H}_2\text{O}$ (Invitrogen 10977015) | as needed to 100.0 $\mu\text{l}$ | |

| Cycler schedule |  |  |
| --- | --- | --- |
| Step | Temperature | Time |
| Digestion | 37°C | 60–90 minutes |
| Pause | 4°C | indefinitely |

Note: Perform a digestion test before performing this step for the first time to validate the time required for the complete digestion of byproducts with your DNA concentration and the chosen restriction enzyme cutting efficiency. The easiest method involves taking the quantity of the original plasmid equal to that used in a recombination reaction (much like in the example in this table), and incubating while taking samples for gel electrophoresis from the reaction volume every 20 minutes. When the gel shows complete digestion, purify the remaining reaction volume and verify on a gel that all of the DNA was digested. In our example using *Ssp*I or *Not*I restriction enzymes, the complete digestion was achieved within 40 minutes for 10 *µ*g of plasmid DNA for all plasmid sizes we used in this study. The digestion takes longer—but not an excessive amount of time—for higher DNA concentrations. For example, for a 300 ng/*µ*l digestion test reaction (30 *µ*g in 100 *µ*l reaction), almost complete digestion of a 13.9 kbp plasmid was achieved in 60 minutes, with no traces of DNA remaining at 90 minutes of digestion (see Supplementary Figure S5).

4. **Use spin column kit to perform final clean up similar to Step 2 above.** Note that the expected mcDNA yield is approximately 30% of the input DNA, and a lower column capacity will suffice. Adjust the number of QIAquick spin columns used accordingly.

For researchers using this protocol for the first time, we recommend using at least 40 *µ*g of source plasmid for the initial experiment. In step 4 above, opt for a low elution volume of 20–25 *µ*l. This will allow to become familiar with the protocol and, if necessary, adjust the production scale accordingly.

The spin column purification steps described above are designed for the easy preparation of mcDNA in quantities ranging from 2 to about 30 *µ*g. If larger quantities are required, scaling up the number of QIAquick columns may become impractical. In such cases, the cleanup steps can be performed more economically using isopropanol precipitation [58] for step 2 of the protocol above. However, lower DNA extraction efficiency should be expected with methods other than modern spin columns. In a comparison study [59], isopropanol precipitation reported DNA recovery of 52.47 ± 19.69% (see also [60]), centrifugal dialysis 12.58 ± 7.15%, and magnetic bead extraction 9.92 ± 3.89%, all notably lower than the above spin column efficiencies.

### Fine-tuning recombination buffer composition for optimal performance

The mcDNA yield depends on the reaction buffer used, the source plasmid size, and a combination of DNA and ΦC31 integrase proportions and concentrations, recombination reaction temperatures, and recombination time. We explored this multivariate space while focusing on determining the optimal conditions for practical mcDNA yield.

The work began with the validation of the buffers facilitating recombination. We tested the buffers used with ΦC31 integrase in other studies [21, 27, 52], as well as a number of buffers customarily used for restriction enzyme reactions sourced from NEB. We found that all NEB buffers were not suitable for the recombination reactions, resulting in all DNA being completely digested in our preliminary tests of the protocol. The buffers containing only NaCl, Tris, and EDTA performed best, resulting in visually complete recombination of the DNA product in 12 hours (see Figure 1d).

We tested whether customarily used additives improve recombination efficiency. Formulations of the buffers with 1 to 5% glycerol had no noticeable difference in the recombination speed. The final recombination reaction mix ultimately contained 1% or more glycerol from the ΦC31 integrase working buffer. NaCl was most effective in the 100–150 mM range, with diminished performance at lower or higher concentrations. We also tested the addition of 100 *µ*g/ml BSA and 5 mM spermidine—as used in [21, 52]—and found that while complicating the handling, these additives had a slight negative effect on the recombination efficiency and thus were not used. Therefore, the final reaction concentrations were 20 mM Tris, 0.1 mM EDTA, 100 mM NaCl (used in the form of the 2X buffer described in the Methods), and quantities of ΦC31 integrase and plasmids DNA as required per experiment. This minimal buffer achieved complete recombination of the DNA product shown in Figure 1d, thus it makes sense that additives beyond this would have no effect at best, and at worst interfere with the reaction.

### DNA purification after the completion of recombination

Initially, we expected to develop a single-tube protocol where the recombination would be followed by the cutting of the bacterial backbone using restriction enzymes and digestion with T5 exonuclease within the same reaction tube, similar to what we successfully achieved for the ligation reaction of CVs (see Oliynyk & Church [35]). This would be particularly helpful because DNA purification spin column yields diminish with increasing DNA sizes. However, we found that the presence of ΦC31 integrase consistently reduced the on-target efficiency of the restriction enzymes we tested (*Not*I, *Ssp*I, and *Psi*I).

Simultaneously, all of these enzymes exhibited star activity, leading to the deterioration of well-recombined mcDNA. We hypothesized that this behavior may be associated with ΦC31 integrase molecules binding with similar affinity to attP and attB, as well as to attL and attR [28, 29], while normally not effecting the reversal of recombination. It is possible that ΦC31 intermittently binds to other plasmid locations, or co-binds in the presence of restriction enzymes targeting specific DNA sequences and modifies their activity. Using the thermal inactivation of ΦC31 integrase at 80°C has not abolished this behavior, and the use of thermolabile Proteinase K [61] with subsequent heat inactivation has not completely resolved the issue, with a fraction of partially digested DNA fragments consistently present in Plasmidsaurus sequencing. After the above testing, it was determined that we could not dispense with the cleanup step after the completion of recombination. Additionally, the proteinase K digestion of ΦC31 integrase after recombination reaction was necessary before the column cleanup, which was also rendered less effective by ΦC31 integrase, with spin columns exhibiting clogging—particularly with higher ΦC31 integrase concentrations. As a result, the protocol included ΦC31 integrase digestion after the completion of recombination using proteinase K, followed by spin column cleanup of the DNA. This was followed by the cutting of the bacterial backbone with a suitable restriction enzyme and the digestion of all non-circular or nicked DNA using T5 exonuclease, followed by final spin column cleanup.

DNA purification was performed using QIAGEN’s *QIAquick PCR Purification Kit* (QIAGEN Cat#28106) on a QIAGEN vacuum manifold and *Eppendorf 5425 R* microcentrifuge, per the manufacturer’s instructions.

### Component concentrations in recombination experiments

As shown in the Results, the mcDNA yield depends on the source plasmid sizes, as well as a combination of DNA and ΦC31 integrase proportions and concentrations, recombination reaction temperatures, and times. A thorough mapping of such multivariate space would take a large number of experiments. Our task was simplified by focusing on finding optimal practical concentrations useful for mcDNA production, which allowed us to rapidly pinpoint the useful reaction proportions. All experiments in Figure 2 were performed using the Protocol steps described earlier, with the following particulars. The concentrations used were (numbers following components are in ng/*µ*l): Figure 2a,b: ΦC31 100, DNA 100; Figure 2c,f: ΦC31 25–200, DNA 100; Figure 2d,g: ΦC31 100, DNA 100–300; Figure 2e,h: ΦC31 100, DNA 100; Figure 2j: ΦC31 200, DNA 66.6.

Figure 2i used all of the same data points displayed in Figure 2c–h, presenting them as molar ratios of ΦC31 integrase molar concentrations relative to DNA molar concentrations. In this figure, ΦC31 integrase concentration was adjusted to account for 85% purity, as reported by GenScript quality analysis (Supplementary Figure S10). The molar ratios were calculated based on ΦC31 integrase containing 611 residues and a corresponding molecular mass of 68 kDa. The molecular masses of corresponding plasmids were calculated based on plasmid size [62], and the molar ratios were calculated as multiples of ΦC31 integrase molecules per DNA molecule for corresponding ΦC31 and DNA concentrations.

### CRISPR-Cas9 base editing: transfection, editing, handling, and sequencing of edited cells

For all base editing validation experiments, we used plasmid *p-att-ef1a-ABE8e-dCas9-BSD* (see also Plasmid design and assembly further below) and mcDNA produced from this plasmid with our proposed protocol. For gRNA expression, we used the CV method described in Oliynyk & Church [44], with a minimal protocol simplification [35]. Specifically, the NEB Ligation Buffer in the ligation step was substituted by NEB rCutSmart buffer (Cat#B6004), in the identical proportion. To compensate for NEB rCutSmart buffer not containing ATP, the ATP amount in the reaction was doubled to achieve the CV protocol original ATP concentration. This simplified the reagents for the digestion step of the CV protocol, allowing, instead of preparing a four-component digestion buffer, just to add 4 *µ*l of T5 Exonuclease and optionally 0.4 *µ*l of additional NEB rCutSmart buffer per 100 *µ*l of CV ligation reaction, with identical final yield ***(this modified CV protocol has passed the peer reviews and is currently in preparation for publication in Nature Protocols, and will be cited here instead of this note in case of NAR publication)***. The CV sequences are available in the Supplementary Data. When needed, mcDNA was further vacuum-concentrated using Thermo Fisher Scientific Savant*^T M^* SpeedVac*^T M^* . Both the HEK293T and mESC lines were transfected using a Neon Transfection System (Thermo Fisher Scientific) using 10*µ*l tips (Cat#MPK1096K). Cells were resuspended at 40–70% confluency, centrifuged at 300G, and resuspended in Neon Buffer R to accommodate 40k cells per edit—similar for both cell lines. At this point, the editing steps differed as follows:

- **Editing and selection of HEK293T cells.** We used 1400 ng of dCas9-ABE8e mcDNA (size 6833 bp) and dCas9-ABE8e plasmid (size: 9533 bp, endotoxins 0.57 EU/*µ*g, Supplementary Figure S19–S20), with 400 ng of the gRNA CVs to achieve a ∼4X molar excess of guide expression vector to a protein expression vector, performing edits in triplicate, following [35]. The transfection was performed using a Neon protocol with two pulses at 1,150V and a duration of 20ms [63]. Edited HEK293T cells were seeded on 24-well plates (Corning) in DMEM supplemented with 10% FBS. Then, 16–18 hours after transfection, 10 *ng/µ*l of blasticidin (Thermo Fisher Scientific) was used to select cells containing the base editor (based on a recommendation by Xiong et al. [64]). Selection continued for 2 days, noting that the control, non-transfected cells died 48 hours into the selection process; at this point the cells were passaged in media with 10 *ng/µ*l of blasticidin. The media was changed with fresh media 24 hours later. Genomic DNA was extracted 120 hours after the beginning of selection using a Zymo Research Quick-DNA Microprep Kit (Cat#D3020). Genomic DNA processing and sequencing were performed similarly for both cell lines, as described further.
- **Editing and selection of mESCs.** The initial transfection was performed with the same amount of DNA as above for HEK cells, with more cell mortality than expected. Thus, the DNA concentration was decreased by approximately four-fold, using 380 ng of dCas9-ABE8e mcDNA or 380 ng of plasmid with 100 ng of gRNA CV. This concentration resulted in the typical cell survival. The transfection was performed using a Neon protocol with two pulses at 1,200V and a duration of 20ms [65]. Edited mESCs were seeded on 24-well plates (Corning) treated with Matrigel (Corning) in ESGRO-2i Medium (Millipore). For mESCs, we tested editing in quintuplicate with mcDNA and in duplicate with plasmid, as presented in the Results. The blasticidin selection was performed 14–16 hours after transfection with dose of 5 *ng/µ*l, and same dose of 5 *ng/µ*l was added 24 hours later and the cell cultures were passaged 48 hours after beginning of selection with fresh media. Genomic DNA was collected two days later in the same manner as for HEK293T cells.
- **Genomic DNA processing and sequencing.** Genomic areas of interest were amplified from 6 ng of whole genome DNA using HiFi Hot Start Readymix (Kapa) at 30 cycles according to the master mix protocol (see Supplementary Table S2 for the list of primers). This was followed by DNA separation via electrophoresis on 1% E-Gel EX Agarose Gels (Invitrogen) and extraction using a Monarch DNA Gel Extraction Kit (T1020S) with an additional washing step. Sanger sequencing was performed in duplicate for each physical sample using Azenta Genewiz with our PCR primers (Supplementary Table S2).

The editing efficiency analysis was performed using EditR [66, 67] is summarized in Figure 4 and illustrated in Supplementary Figures S6–S7.

### HITI editing: transfection, editing, handling, and sequencing of edited cells

- **Transfection for HITI experiments.** HEK293T cells were seeded into 24-well plates at a density of 120K per well 20-24 hours before transfection. At approximately 70% confluency, cells were transfected using Lipofectamine 3000 (Thermo Fisher Scientific) following the manufacturer’s instructions. Briefly, 250 ng of Cas9 plasmid (pCAG-1BPNLS-Cas9-1BPNLS, Addgene #87108), 200 ng of gRNA CVs, and 300 ng of HITI mcDNA template were mixed in 50 *µ*l of Opti-MEM with 1 *µ*l of 3000 Reagent and 1.5 *µ*l of Lipofectamine 3000 Reagent per well. 48 hours later, cells were harvested by removing the media, washed one time with PBS, and then trypsinized using TrypLE Express (Thermo Fisher Scientific, Cat#12604013) for 3 minutes. Resuspended cells were pelleted via centrifugation for 5 minutes at 500 g. After centrifugation, cells were resuspended in media containing hygromycin (Thermo Fisher Scientific, Cat#10687010) at a concentration of 50*µ*g/ml and seeded in 6-well plates. Cells were allowed to grow for 3 days, then the hygromycin concentration was increased to 200 *µ*g/ml. Continuous hygromycin treatment was carried out for 14 days until all unedited cells were eradicated, as measured by observing a control well with untransfected cells and treatment with the same hygromycin concentration.
- **Flow cytometry analysis.** Hygromycin-selected cells and untransfected control cells were harvested as described above. After centrifugation, cell pellets were resuspended into PBS with 2% FBS and 1 *µ*g/ml of 4’,6-diamidino-2-phenylindol (DAPI) (Thermo Fisher Scientific, Cat#D1306) and transferred into 5 ml round bottom polystyrene flow tubes with cell strainer (Corning Life Sciences, Cat#352235). Cells were analyzed by flow cytometry (Symphony A1, BD Biosciences) and data was processed with FlowJoTM 10 software. An overview of gating and analysis is given in Supplementary Figure S8.
- **Genomic DNA extraction and PCR analysis.** To confirm the correct integration of HITI templates, genomic DNA was extracted from HITI-treated cells and untransfected cells (as negative controls) using a Zymo Research Quick-DNA Microprep Kit (Cat#D3020). Genomic DNA processing and sequencing were performed as described above with slight modifications. ACTB or GAPDH-specific forward primers were used with a common mKate reverse primer to amplify the genomic DNA region spanning the junction of the HITI-mediated integration. Q5 Hot Start High-Fidelity 2X Master Mix (NEB., Cat#M0494) was used to amplify the genomic region using 30 cycles of amplification with 20ng of total genomic DNA per reaction. PCR amplicons were cleaned up using QIAGEN’s QIAquick PCR Purification Kit (QIAquick) and separated by electrophoresis on 1% E-Gel EX Agarose Gels (Invitrogen). For sequencing, the Monarch DNA Gel Extraction Kit (T1020S) was used to purify the excised DNA band from the gel and sent for Sanger sequencing. Sanger sequencing was performed using Azenta Genewiz with a primer specific to mKate insert. All primers are listed in Supplementary Table S2.

#### Plasmid design and assembly

We used *p-dCas9-SSAP-MS2-BB* (Addgene Cat#183826) [68] as a convenient source of bacterial backbone and dCas9 for Gibson assembly. The insert of the BSD gene is based on our earlier publication [44], while the 10482 bp plasmid/mcDNA (named *p-att-ef1a-MS2-RecT-dCas9-BSD*, available at Addgene #226108) is used in our other ongoing project, and our dCas9-ABE8e plasmid (named *p-att-ef1a-ABE8e-dCas9-BSD*) is a modification of this plasmid with TADa for ABE base editing derived from [11] and cloned using Gibson assembly NEBuilder HiFi DNA Assembly Master Mix (NEB Cat#E2621) and amplified using NEB 5-alpha F‘Iq Competent *E. coli* (NEB Cat#C2992H). The necessary dsDNA fragments were ordered from Twist Bioscience.

The plasmid *p-att-ef1a-MS2-RecT-dCas9-BSD*, 10482 bp in length, is a functional plasmid that was used as a basis for the recombination experiments (available at Addgene #226116). Two modifications of this plasmid—a truncation to create a 5758 bp plasmid *p-att-ef1a-MS2-RecT-dCas9-BSD-truncated* and an extension to create a 13942 bp plasmid *p-att-ef1a-MS2-RecT-dCas9-BSD-extended*—were used for recombination experiments but are not intended to be functional for genome editing. An additional even shorter 3871 bp parental plasmid *p-att-SuperShort1211bp*, designed to test the yield of 1211 bp mcDNA was produced from *p-att-ef1a-MS2-RecT-dCas9-BSD-truncated* plasmid by cutting it at *Age*I and *Bbs*I restriction sites and inserting a short DNA sequence to produce desired mcDNA length; similar to *p-att-ef1a-MS2-RecT-dCas9-BSD-truncated*, this plasmid is also designed exclusively for recombination testing purposes, and does not contain a valid protein expression vectors. The sequences for all aforementioned plasmids are available in the *Plasmids sequences* folder in the Supplementary Data.

#### Testing of endotoxin levels

To test the endotoxin levels for the source plasmid and the resulting mcDNA we used Charles River Endosafe® nexgen-MCS endotoxin testing equipment and PTS2001F Limulus Amebocyte Lysate (LAL) endotoxin testing cartridges certified for the FDA validation of endotoxin levels, helped by the expertise of the Wyss Institute at Harvard University that routinely performs such tests for biological materials intended for FDA validation. Per recommendation of Charles River technical support we used 10% dilution of Charles River BD100 dispersant, which is critical for reduction of the clumping and sticking of endotoxins in DNA samples, instrumental in achieving consistent and reliable testing results. The PTS cartridges contain 4 lanes each for every individual sample, two for spike calibration and two for the endotoxin level measurement and cross-validation, resulting in highly accurate measurements.

For initial tuning, we used an earlier prepared with the use of QIAGEN Midiprep Plus *p-att-ef1a-ABE8edCas9-BSD* plasmid and resulting mcDNA produced with the help of Plasmid2MC. We found the source plasmid endotoxin level at 1.88 EU/*µ*g and the Plasmid2MC produced mcDNA with 0.029EU/*µ*g. This presented a 60-fold improvement in endotoxing levels (see the reports Supplementary Figure S15). For this test we used the FDA approved cartridgec Cat#PTS2005F with the endotoxin range of 5–0.05 EU/ml, using dilution factor 2:1. This dilution allowed us to achieve an accurate reading for the mcDNA, however it showed that with such low endotoxin levels, the aforementioned cartridges PTS2001F with 1–0.01 EU/ml range were more appropriate, and we used them in all subsequent tests. The DNA was prepared in stock 1:1 concentrations of 1 *µ*g DNA per ml, and the further necessary dilutions were performed as required per Charles River instructions. Endotoxin level tests were performed in triplicate.

## Supplementary information

Supplementary information accompanying this manuscript is available in a file Plasmid2MCsupplementary.PDF.

## Data availability

The raw data for the results presented in this study can be found in a single SupplementaryData.ZIP file, composed of the following items:

1. *Recombination Yields Table.xlsx*.
2. *Base Editing Efficiency Sanger-EditR Results.xlsx*.
3. Base Editing plasmid, mcDNA and gDNA CV sequences.
4. HITI plasmid, mcDNA and gDNA CV sequences.
5. Sequenced Plasmidsaurus recombined mcDNA.

Plasmids: *p-att-ef1a-MS2-RecT-dCas9-BSD* is deposited at Addgene #226108, and *p-att-ef1a-ABE8e-dCas9-BSD* is deposited at Addgene #226116.

## Acknowledgments

A.M. was supported by the Ibn Rushd Postdoctoral Fellowship from King Abdullah University of Science and Technology. E.K. received funding through SRA with Colossal Biosciences. We thank Shanda Lightbown from the Wyss Institute at Harvard University for providing access to equipment and assistance with endotoxin testing.

## Author contributions statement

R.O. designed this research and performed Plasmid2MC and base editing experiments; R.O. and A.M. collaborated on conceptual foundations; E.K. defined mouse genomic targets for mESC base editing; A.M. conceived and performed HITI experiments and analysis; R.O., A.M. and G.C. analyzed results and wrote the manuscript.

