## Supplementary Information for "Plasmid2MC: Efficient cell-free recombination of plasmids into high-purity minicircle DNA for use in genome editing applications"

**Roman Teo Oliynyk,[1,2,*] Ahmed Mahas,[1] Emil Karpinski[1] and George M. Church[1,3]**

[1]Department of Genetics, Harvard Medical School, Boston, 02115, MA, USA, [2]Department of Computer Science, University of Auckland, Auckland, 1010, New Zealand and [3] Wyss Institute for Biologically Inspired Engineering at Harvard University, Boston, 02115, MA, USA

#### Contents

Reagents and resources 2
Unprocessed gel photo for Figure 1 4
mcDNA sequencing histograms 5
Gel electrophoresis image of dCas9-ABE8e mcDNA and three gRNA CVs 6
Digestion test for Protocol Step 3 7
Sanger sequencing - EditR figures 9
Base editing guides and primers 11
HITI flow cytometry 12
HITI gel electrophoresis images and analysis 13
ΦC31 amino acid sequence 14
Comparative evaluation of System Biosciences Inc MC-Easy™ Minicircle DNA Production Kit 15
The collection of the Charles River nexgen-MCS endotoxin test reports 22
Recombination efficiency as a function of ΦC31 molecules per kbp of plasmid DNA 26

#### Reagents and resources

**Supplementary Table S1.**

| Reagent or resource | Source | Cat# |
|---|---|---|
| SspI-HF | New England BioLabs | R3132L |
| NotI-HF | New England BioLabs | R3189L |
| AgeI-HF | New England BioLabs | R3552S |
| FspI | New England BioLabs | R0135S |
| T5 Exonuclease | New England BioLabs | M0663L |
| Thermolabile Proteinase K | New England BioLabs | P8111S |
| Proteinase K | New England BioLabs | P8107S |
| TrypLE | Thermo Fisher Scientific | 12605010 |
| Puromycin Dihydrochloride | Thermo Fisher Scientific | A1113803 |
| Blasticidin S HCl | Thermo Fisher Scientific | A1113903 |
| Penicillin-Streptomycin | Thermo Fisher Scientific | 15070063 |
| Gibco DMEM, high glucose, pyruvate | Thermo Fisher Scientific | 11995040 |
| ESGRO-2i Medium | Millipore | SF016-100 |
| Embryonic Stem Cell FBS | Thermo Fisher Scientific | 16141002 |
| Nuclease-free H2O | Invitrogen | 10977015 |
| HiFi Hot Start - Readymix | KAPA Biosystems | KK2602 |
| Q5 High-Fidelity Master Mix | New England BioLabs | M0492L |
| NEBNext Ultra II Q5 Master Mix | New England BioLabs | M0544L |
| QIAquick PCR Purification Kit | QIAGEN | 28106 |
| Sodium Acetate Solution (3 M), pH 5.2 | Thermo Fisher Scientific | R1181 |
| Monarch PCR & DNA CleanUp Kit | New England BioLabs | T1030S |
| Monarch DNA Gel Extraction Kit | New England BioLabs | T1020S |
| NucleoSpin Gel and PCR Cleanup | Macherey-Nagel | 74609.50 |
| MC-Easy Minicircle DNA Production Kit | System Biosciences | MN920A-1 |
| QIAGEN Plasmid Plus Midi Kit | QIAGEN | 12943 |
| QIAGEN Plasmid Maxi Kit | QIAGEN | 12163 |
| QIAGEN EndoFree Plasmid Maxi Kit | QIAGEN | 12362 |
| Invitrogen 1% E-GELS EX | Invitrogen Corporation | G402021 |
| NEB 5-alpha F'Iq Competent *E. coli* | New England BioLabs | C2992H |
| HEK293T | ATCC | CRL-3216 |
| ESGRO Adapted C57/BL6 Mouse ESC | Millipore | SF-CMTI-2 |
| p-dCas9-SSAP-MS2-BB | Addgene | 183826 |

Continued on next page

**Supplementary Table S1.** (Continued)

| Reagent or resource | Source | Cat# |
|---|---|---|
| NEBuilder HiFi DNA Assembly Master Mix | New England BioLabs | E2621 |
| Tris-EDTA, pH 8.0 | Fisher Scientific | BP2473 |
| 1M Tris-HCl, pH 8.0 | Corning | 46-031-CM |
| Matrigel | Corning | 354277 |
| Lipofectamine 3000 | Thermo Fisher Scientific | L3000015 |
| Opti-MEM™ Reduced Serum Medium | Thermo Fisher Scientific | 31985062 |
| PBS, pH 7.4 | Thermo Fisher Scientific | 10010023 |
| Hygromycin B (50 mg/mL) | Thermo Fisher Scientific | 10687010 |
| FBS, Premium, heat-inactivated | Thermo Fisher Scientific | A5670801 |
| DAPI | Thermo Fisher Scientific | D1306 |
| TrypLE Express | Thermo Fisher Scientific | 12604013 |
| Q5 Hot Start High-Fidelity 2X Master Mix | New England BioLabs | M0494 |
| 5 ml round bottom polystyrene flow tubes | Corning Life Sciences | 352235 |
| pCAG-1BPNLS-Cas9-1BPNLS | Addgene | 87108 |
| Quick-DNA Microprep Kit | Zymo Research | D3020 |
| QIAvac 24 Plus vacuum manifold | QIAGEN | 19413 |
| Eppendorf 5425 R microcentrifuge | Eppendorf | 5406000240 |
| NanoDrop Eight Spectrophotometer | Thermo Fisher Scientific | NDE-GL |
| mySPIN® 6 Mini Centrifuge | Thermo Fisher Scientific | 75004061 |
| NanoDrop™ 8000 Spectrophotometer | Thermo Fisher Scientific | ND-8000-GL |
| Eppendorf® ThermoMixer heat block | Eppendorf | 5382000023 |
| Eppendorf® Mastercycler® Pro | Eppendorf | 950030010 |
| Endosafe® nexgen-MCS™ package | Charles River | MCS650K |
| Endosafe® LAL Cartridge, 1–0.01EU/ml, FDA | Charles River | PTS2001F |
| Bio-Dispersing Agent BD100 | Charles River | BD100 |

#### Unprocessed gel photo for Figure 1

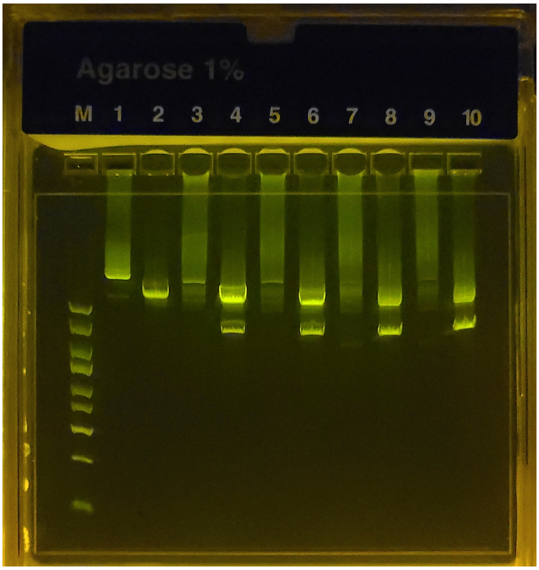


**Supplementary Figure S1**: **Gel image showing the ΦC31 recombination of 10418bp plasmid into a 2.7kbp bacterial backbone circle and 7758bp DNA circle (unprocessed gel photo).** Each lane contains 300ng of DNA.

L1 - pure supercoiled plasmid; L2 - the same plasmid with a single cut with a restriction enzyme (NotI); here, linear DNA travels faster than supercoiled DNA, and L1 presents a very weak matching band, showing a minuscule amount of linear plasmid.

L3 - after 80 minutes of recombination showing diffuse DNA scattering corresponding to the varying speed of travel of miscellaneous catemers depicted in main article Fig. 1b. L4 - the same product cut with NotI, which is present in a single location on the bacterial backbone, thus allowing bacterial backbone release from the tangle, resulting in distinct bands (2.7 and 7.7kbp) and a weaker band corresponding to the non-recombined plasmid, NaCl 100mM. L5, L6 - same as above with NaCl 150mM buffer;

L7 and L8 - same as L3 and L4 after 12 hours of recombination; the original plasmid band is now almost invisible, NaCl 100mM. L9 and L10 - same as L7 and L8 with NaCl 150mM buffer.

It can be seen that the ΦC31 recombination reactions proceeded similarly between the 100 and 150mM NaCl buffer concentrations, resulting in similar gel results, with the 150mM reaction minimally visually lagging after 12 hours of reaction.

#### mcDNA sequencing histograms

In every case, the Plasmidsaurus long read sequencing resulted in a perfect match of the expected and produced mcDNA recombining across attP and attB sites. The sequencing files can be seen in the Supplementary Data. Shown here are samples of the recombination results for three tested source plasmids and mcDNA sizes, with source plasmid DNA concentrations of 100 and 300ng/$\mu$l. Intramolecular recombination was a preferred outcome for both DNA concentrations, with only a small fraction of recombinations resulting in the dimer concatemer at both plasmid concentrations.

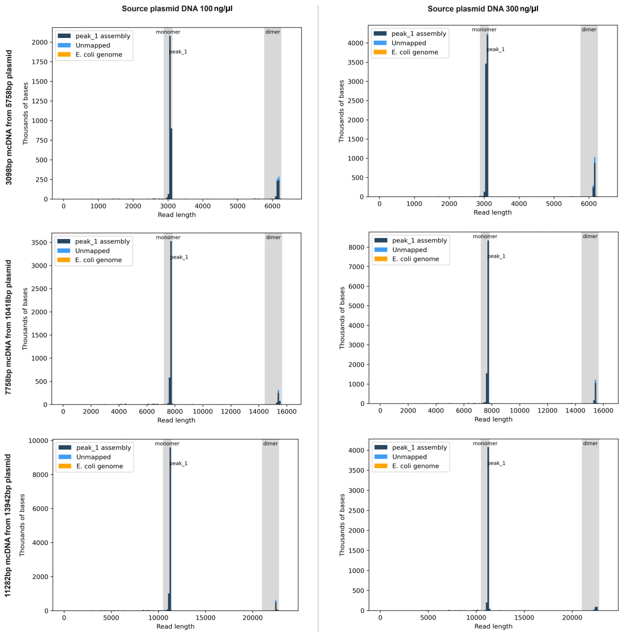


**Supplementary Figure S2**: **Sequencing diagrams produced by Plasmidsaurus** The y-axis numbers represent the total number of bases read, which are merely relative numbers for our purposes; a higher number of reads corresponds to better assembly quality and fewer spurious unmapped blocks. The x-axis is the mcDNA assembled read length. The proportion of dimers relative to the intended mcDNA, while slightly higher for plasmid DNA concentrations of 300ng/$\mu$l, represented only a minor fraction at all concentrations.

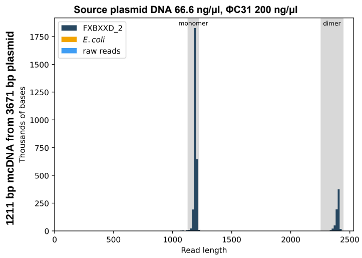


**Supplementary Figure S3**: **Sequencing diagrams of 1211bp mcDNA for the shortest mcDNA tested.** Produced by Plasmidsaurus. The y-axis numbers represent the total number of bases read, which are merely relative numbers for our purposes; a higher number of reads corresponds to better assembly quality and fewer spurious unmapped blocks. The x-axis is the mcDNA assembled read length.

#### Gel electrophoresis image of dCas9-ABE8e mcDNA and three gRNA CVs

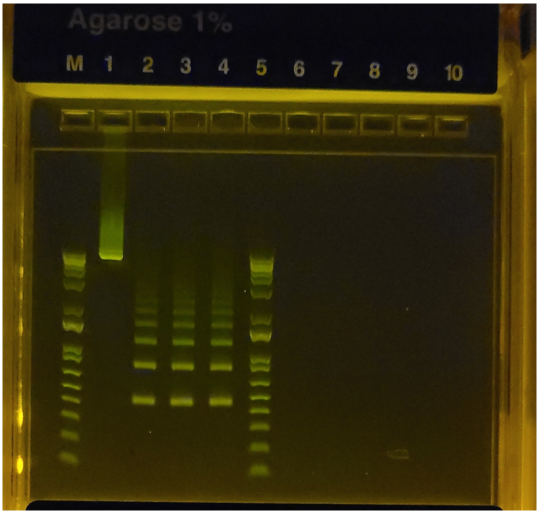


**Supplementary Figure S4**: **Gel electrophoresis image of dCas9-ABE8e mcDNA and three gRNA CVs.** Lanes: L1 - a 6873bp dCas9-ABE8e mcDNA, recombined from a 9533bp source plasmid; L2–L4 - three 457bp CVs for the expression of gRNAs for validation editing in HEK293T cells, showing the primary circle and the second and higher-order concatemers, all of which are effective as gRNA expression vectors. (Unprocessed image).

#### Digestion test for Protocol Step 3

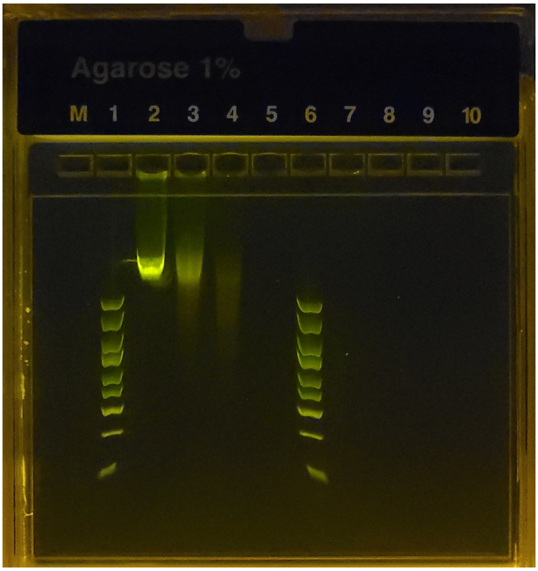


**Supplementary Figure S5**: **Digestion test for Step 3 of the recombination protocol.** Lanes: L1 and L6 - DNA ladder; L2 - 20-minute digestion (2 $\mu$l sample); L3 - 40-minute digestion (2 $\mu$l sample); L4 - 60-minute digestion (2 $\mu$l sample); L9 - 90-minute digestion (purified remaining reaction). (Unprocessed image).

Perform this digestion test before performing the digestion step in the minicircle recombination protocol for the first time to determine the time required for the complete digestion of byproducts with your DNA concentration and restriction enzyme cutting efficiency. The easiest way involves taking the same quantity of the original plasmid—as per the example in Table—and incubating while taking samples for gel electrophoresis from the reaction volume every 20 minutes. When the gel shows complete digestion, purify the remaining reaction volume and verify on a gel that all DNA was digested. In our example using SspI and NotI, complete digestion was achieved within 40 minutes for 10$\mu$g of plasmid DNA for all plasmid sizes. The digestion takes longer—but not an excessive amount of time—for higher DNA concentrations. It is important to scale the buffers to maintain the proportions of the restriction enzyme(s) and T5 exonuclease for fast reaction times.

The example in Supplementary Figure S5 shows the digestion test reaction at a DNA concentration of 300ng/$\mu$l (30$\mu$g in a 100$\mu$l reaction). Here, the DNA concentration before the start of digestion was 300ng/$\mu$l. Then, 2$\mu$l were sampled every 20 minutes. Thus, the pre-digestion DNA amount in 2$\mu$l equals 600ng per lane. Incrementally more complete digestion was observed at 20 and 40 minutes, with little DNA remaining after 60 minutes. Lane L5 shows complete digestion when using the purified DNA product of the remaining 92$\mu$l of reaction after 90 minutes of digestion, with no traces of DNA remaining.

#### Sanger sequencing - EditR figures

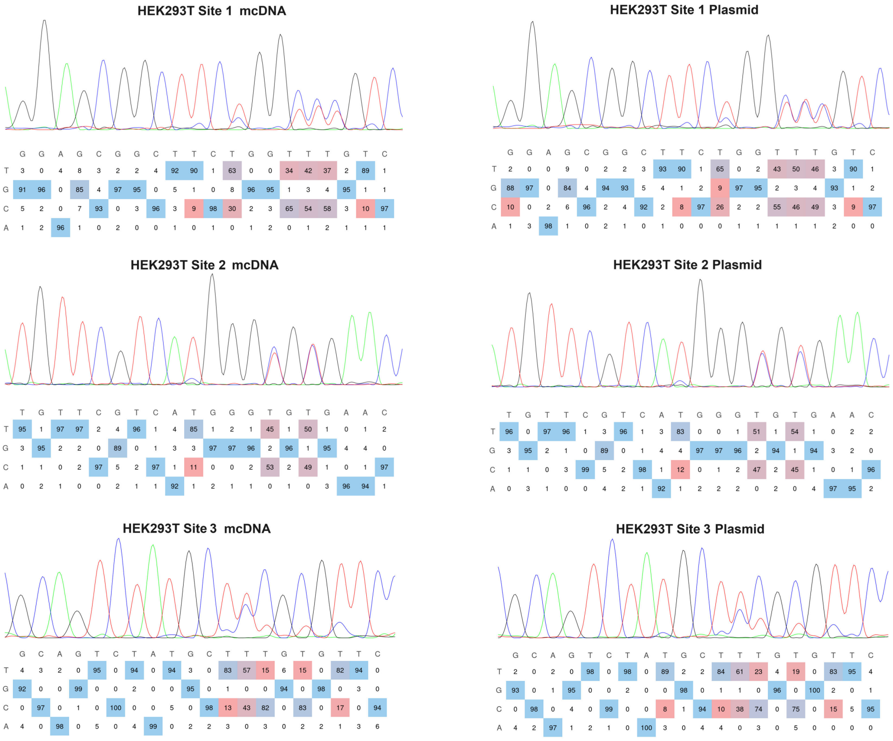


**Supplementary Figure S6**: **Sanger sequencing of dCas9-ABE8e editing HEK293T sites.** Images for three edited sites, using mcDNA and the source plasmid. (EditR analysis).

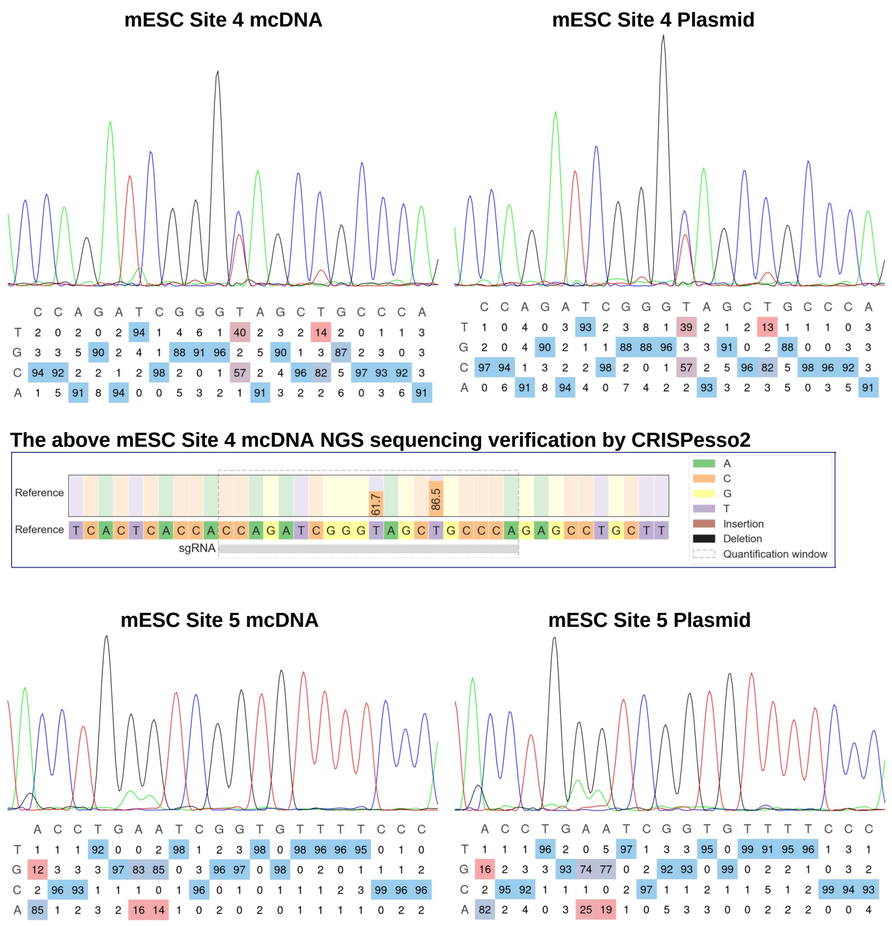


**Supplementary Figure S7**: **EditR analysis of Sanger sequencing of dCas9-ABE8e editing for mESC sites.** Images for the two edited sites, using the mcDNA and the source plasmid. CRISPResso2 [1] Nucleotide distribution around the sgRNA TGGGCAGCTACCCGATCTGG. for data point sample 3 in mESC site 4 confirms the Sanger sequencing results of editing efficiency demonstrated in the Figure 4(C) in the main article. Specifically, for this sample EditR from Sanger sequencing showed 82.89% and 56.84%, with NGS finding 86.5% and 61.7% respectively; as expected, the Sanger sequencing noise would statistically attribute 3/4 negative vs 1/4 positive noise bias. Thus, true editing efficiency values by NGS are slightly higher than the Sanger sequencing derived values.

#### Base editing guides and primers

**Supplementary Table S2.** Guides and primers used in base editing experiments.

| | |
|---|---|
| **HEK293T - Site 1** (chr3, CTNNB1) | |
| Guide **(PAM)** | GACAAACCAGAAGCCGCTCC**(TGG)** |
| Primer FWD | GAACTGGACAAACTTCTAACAAAAGGTATTTGC |
| Primer FWD-I | TTCTTGTAGCCCTCTTTTTATTGGA |
| Primer REV | GTCCACTTACCTATCACAATCACAACTGC |
| **HEK293T - Site 2** (chr12, GAPDH) | |
| Guide **(PAM)** | GTTCACACCCATGACGAACA**(TGG)** |
| Primer FWD | GCTGACTCAGCCCTGCAAAG |
| Primer REV | GCAAAGAAAGAGGGAGCGGG |
| **HEK293T - Site 3** (chr5, VISTA enhancer hs267) | |
| Guide **(PAM)** | GAACACAAAGCATAGACTGC**(GGG)** |
| Primer FWD | CTCAGTGGCAGGACGTCTGC |
| Primer REV | CCAGCCCCATCTGTCAAAC |
| **mESC - Site 4** (chr11, Wnt3a) | |
| Guide **(PAM)** | TGGGCAGCTACCCGATCTGG**(TGG)** |
| Primer FWD | GCCCAAACAGGTCCAAGTGG |
| Primer FWD-I | CAGCCAAGGAAAACAACCCG |
| Primer REV | CGATGGCTCCTCTCGGATACC |
| **mESC - Site 5** (chr2, Prkra) | |
| Guide **(PAM)** | ACCTGAATCGGTGTTTTCCC**(AGG)** |
| Primer FWD | AAGTTATGTCACCAACGGTTACTCTGAAGG |
| Primer FWD-I | TCTGAAGGTGAAAGTGGGCACG |
| Primer REV | GCTTTTGCTTCCCTTTCAGCTTTGACG |
| **HITI: gel image in Fig. 4** | |
| Primer GAPDH-HITI-2-F | TGTGGCTGGGGCCAGAGACTGG |
| Primer ACTB-HITI-2-F | TCACATCCAGGGTCCTCACTGCC |
| Primer mKate-R-2 | CCTGGGTAGCGGTCAGCACGC |
| Primer mKate REV-I | GGTGACTCTCTCCCATGTGAAG |
| **HITI: Supplementary gel image** | |
| Primer ACTB-HITI-F | GAGCTGACCTGGGCAGGTCGG |
| Primer ACTB-HITI-R | GGATGCTCGCTCCAACCGACTG |
| Primer GAPDH-HITI-F | GCCCTCAACGACCACTTTGTCAAGC |
| Primer GAPDH-HITI-R | GATGGTACATGACAAGGTGCGGC |

The inner primers (-I) and/or gel purification were used to achieve better amplicon selection and correspondingly quality of Sanger sequencing.

#### HITI flow cytometry

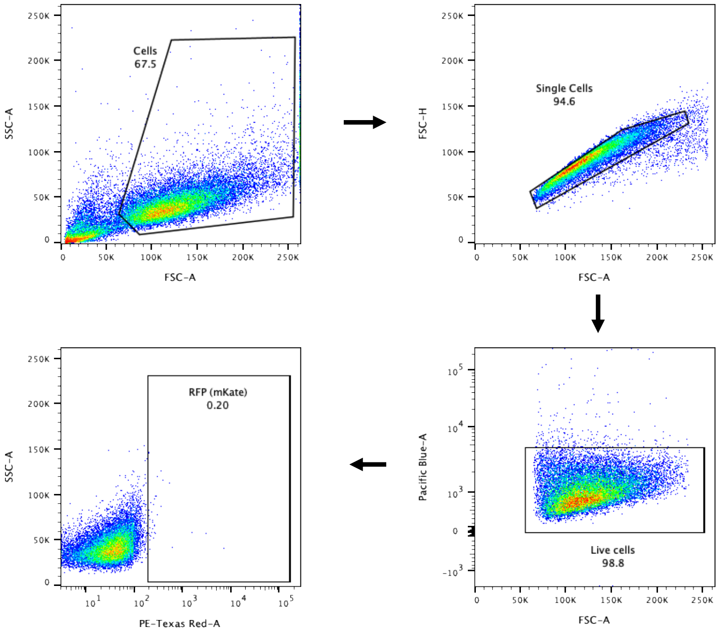


**Supplementary Figure S8**: **Gating strategy used for flow cytometry analysis of HITI experiment in HEK293T cells.** Gates are set using a control population not treated with HITI.

#### HITI gel electrophoresis images and analysis

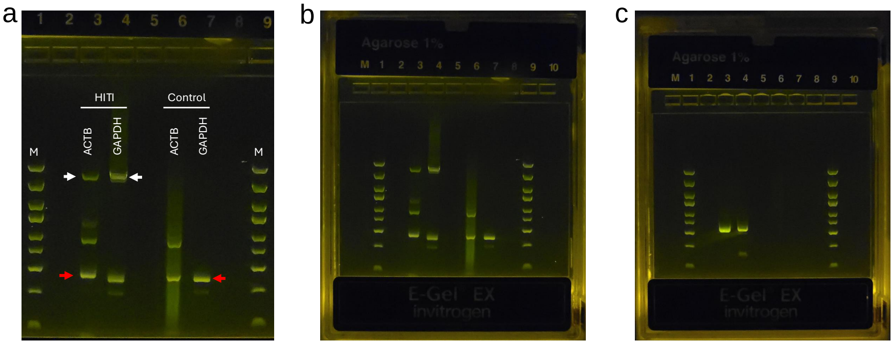


**Supplementary Figure S9**: **HITI gel electrophoresis images.** **(a)** Gel electrophoresis image displaying the PCR amplification results of the entire HITI-integrated donor sequence in targeted genes. Primers flanking the integration site were utilized for amplification. Red arrows indicate the PCR amplicons of the targeted sequences free of HITI integration in both control and HITI samples. The ACTB HITI-free amplicon is 426bp in length, while the GAPDH HITI-free amplicon is 386 bp. White arrows point to the PCR amplicons of the targeted sequences with HITI integration. The ACTB amplicon with HITI integration is 3550bp long, and the GAPDH amplicon with HITI integration is 3511 bp. This gel image illustrates the presence of both HITI-integrated and non-integrated sequences, confirming the integration events in the targeted genes. M: Ladder.
**(b)** Unprocessed image (a). **(c)** Unprocessed image of Fig. 4 in the main text.

#### ΦC31 amino acid sequence

MDTYAGAYDRQSRERENSSAASPATQRSANEDKAADLQREV
ERDGGRFRFVGHFSEAPGTSAFGTAERPEFERILNECRAGRL
NMIIVYDVSRFSRLKVMDAIPIVSELLALGVTIVSTQEGVFRQ
GNVMDLIHLIMRLDASHKESSLKSAKILDTKNLQRELGGYVG
GKAPYGFELVSETKEITRNGRMVNVVINKLAHSTTPLTGPFE
FEPDVIRWWWREIKTHKHLPFKPGSQAAIHPGSITGLCKRMD
ADAVPTRGETIGKKTASSAWDPATVMRILRDPRIAGFAAEVI
YKKKPDGTPTTKIEGYRIQRDPITLRPVELDCGPIIEPAEWYE
LQAWLDGRGRGKGLSRGQAILSAMDKLYCECGAVMTSKRGE
ESIKDSYRCRRRKVVDPSAPGQHEGTCNVSMAALDKFVAERI
FNKIRHAEGDEETLALLWEAARRFGKLTEAPEKSGERANLVA
ERADALNALEELYEDRAAGAYDGPVGRKHFRKQQAALTLRQ
QGAEERLAELEAAEAPKLPLDQWFPEDADADPTGPKSWWG
RASVDDKRVFVGLFVDKIVVTKSTTGRGQGTPIEKRASITWA
KPPTDDDEDDAQDGTEDVAAHHHHHH*

##### Certificate of Analysis

| | |
|---|---|
| **Name:** | phiC31 |
| **Type:** | Recombinant enzyme |
| **Shipping Condition:** | Dry Ice |
| **Lot Number:** | U348F486G0-4/P02JA001 |
| **Expression Host :** | E.coli |
| **Purification:** | Protein was obtained from supernatant of cell lysate, Ni column |
| **Total Amount:** | 18.96 mg. 1.00 ml/tube, 4 tubes |
| **Concentration:** | 4.74 mg/ml(by Bradford) A260/A280: 0.557 |
| **Purity:** | ≥85%(SDS-PAGE under reducing condition) |
| **Endotoxin:** | ≤0.4 EU/mg |
| **Storage And Handling:** | Store at -80°C. Avoid repeated freezing and thawing |
| **Storage Buffer:** | 50 mM Tris-HCl, 500 mM NaCl, 10 % Glycerol, pH 8.0 |

**SDS-PAGE & Western blot Analysis:**

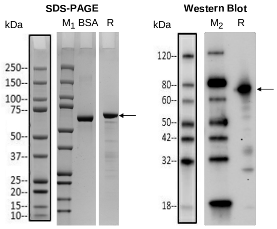


Lane M1: Protein Marker, Bio-rad, Cat. No. 1610374S, refer to annotated key on the left for size
Lane M2: Protein Marker, GenScript, Cat. No. M00673, refer to annotated key on the left for size
BSA: 2.00 μg
R:Reducing condition
Primary antibody: Mouse-anti-His mAb (GenScript, Cat.No. A00186)

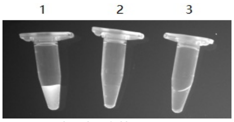

**DNAse detection of phiC31**
Tube 1: Positive Control
Tube 2: Negative Control
Tube 3: phiC31

**Appendix:**

**U348F486G0-1_phiC31**

**Protein Length=611 aa His_6: 606-611**

**MDTYAGAYDRQSRERENSSAASPATQRSANEDKAADLQREVERDGGRFRFVGHFSEAPGTSAFGTAERPEFERILN
ECRAGRLNMIIVYDVSRFSRLKVMDAIPIVSELLALGVTIVSTQEGVFRQGNVMDLIHLIMRLDASHKESSLKSAKILDTK
NLQRELGGYVGGKAPYGFELVSETKEITRNGRMVNVVINKLAHSTTPLTGPFEFEPDVIRWWWREIKTHKHLPFKPGS
QAAIHPGSITGLCKRMDADAVPTRGETIGKKTASSAWDPATVMRILRDPRIAGFAAEVIYKKKPDGTPTTKIEGYRIQRD
PITLRPVELDCGPIIEPAEWYELQAWLDGRGRGKGLSRGQAILSAMDKLYCECGAVMTSKRGEESIKDSYRCRRRKVV
DPSAPGQHEGTCNVSMAALDKFVAERIFNKIRHAEGDEETLALLWEAARRFGKLTEAPEKSGERANLVAERADALNA
LEELYEDRAAGAYDGPVGRKHFRKQQAALTLRQQGAEERLAELEAAEAPKLPLDQWFPEDADADPTGPKSWWGRA
SVDDKRVFVGLFVDKIVVTKSTTGRGQGTPIEKRASITWAKPPTDDDEDDAQDGTEDVAAHHHHHH***

**Supplementary Figure S10**: **ΦC31 integrase protein Certificates of Analysis**

#### Comparative evaluation of System Biosciences Inc MC-Easy™ Minicircle DNA Production Kit

##### Introduction

Our initial foray into producing minicircle DNA (mcDNA) involved using the System Biosciences Inc. (SBI) MC-Easy$^{TM}$ Minicircle DNA Production Kit, hereafter referred to as the MC-Easy$^{TM}$ Kit. At first glance, it appeared to be a solution for our need to produce mcDNA for genome editing applications, despite the kit's high cost: $1,072 for 5 preparations and $1,914 for 10 preparations.

Our initial experience with the MC-Easy$^{TM}$ kit revealed the complexities of working with it. On our first attempt, we failed to achieve the proper optical density and pH, necessitating a restart of the process (for reference on steps, see the MC-Easy$^{TM}$ Kit and manual [2, 3]). We fine-tuned the preparation of the required 400 ml of the kit's ZYCY10P3S2T *E. coli* transformed with the desired plasmid containing our mcDNA expression vector inserted into the MC-Easy$^{TM}$ proprietary bacterial backbone. However, the yield of the mcDNA was typically not much above 100 $\mu$g. We found that the mcDNA produced by the kit, even when showing a perfect gel image, was toxic to sensitive cells. To increase cell viability, we had to reduce the transfected DNA amount to below 100 ng per edit using Lonza and Neon editing systems, at which point editing efficiency was only negligible.

Applying the recommended additional restriction enzyme cutting of the bacterial backbone and the parental plasmid, followed by the ATP-dependent DNase reagent provided by SBI, digesting all DNA except the desired mcDNA, and the subsequent precipitation and cleanup reduced the quantity of remaining mcDNA. However, the cell toxicity persisted. Reviewing the literature, we found that cleanup of mcDNA produced by such kits is challenging and often results in mcDNA loss [4, 5, 6, 7]. We suspected that high endotoxin levels might be the problem; however, like our colleagues, we had neither the experience nor the need to test endotoxin levels when preparing plasmids.

In need of a reliable method for preparation of high quality mcDNA, we developed our Plasmid2MC method, which has proven remarkably effective. During the preparation of this publication, the reviewers requested a systematic evaluation of the MC-Easy$^{TM}$ kit's minicircle product, particularly regarding its endotoxin levels.

##### Difficulties using System Biosciences MC-Easy™ Kit

Here we first summarize very time- and labor- consuming steps involved with mcDNA preparation using SBI MC-Easy™ Kit, and brief summary of the endotoxin results that will be further explored in the next section. The SBI MC-Easy™ Minicircle DNA Production Kit Manual [3] describes a multi-day, time and labor-consuming, expensive procedure with many failure points. We recommend reviewing the original manual for more detail, but here is an abbreviated summary of the MC-Easy™ Kit, followed by a comparison with the much simpler, more efficient and reliable Plasmid2MC method. To ensure a fair comparison, we are assuming that preliminary steps of plasmid assembly and cloning are identical for both methods, and starting at the point when the source plasmid (also called parental plasmid) is validated, plated, and has produced bacterial colonies ready to use.

**MC-Easy™ Kit - time-consuming, expensive steps with many failure points:**

- **Day 1 per SBI:** A single colony is inoculated in 2 ml of LB containing 50 $\mu$g/mL kanamycin, incubated at 30°C shaking at 250 rpm for 4-6 hours. Then, 0.5-1 ml of the resulting bacterial culture is inoculated into 200 ml of 1X growth medium and returned to the incubator at 30°C shaking at 250 rpm overnight.
- **Day 2 per SBI:** After overnight growth (in our experience, starting OD600 testing early at 12 hours as OD600 may increase faster than expected, requiring restarting the production as described below), the pH and OD600 of the culture medium need to be measured. The pH should be around 7, and the OD600 should be in the range of 4–6.
  - **Failure Point 1** occurs if OD600 exceeds 8 or pH drops below 6.5, in such case SBI requires restarting the protocol from the beginning.
  - If OD600 is in the 4-6 range, combine with 200 ml of 1X Induction medium for a total volume of 400 ml. If OD600 is 6-8 with pH >6.5, recovery is still possible, but 400 ml of Induction medium must be used, leading to a 600 ml culture volume, which complicates extraction and requires eventual extra Induction medium purchase from SBI since the kits come with equal volumes of growth and induction media.
  - The combined mixture is then incubated at 30°C, shaking at 250 rpm for 3 hours, followed by increasing the temperature to 37°C for 1 hour, totaling 4 hours; longer induction increases bacterial death and contamination (**Failure Point 2**, requiring restarting the protocol).
  - Performing a miniprep on 1 ml of culture, followed by restriction digest for quality check, while the remaining culture volume is centrifuged, pelleted and pellet stored at -20°C or -80°C. If quality is confirmed (if it is not, **Failure Point 3**, requiring restarting the protocol), proceed with Maxiprep recommended by SBI.
- **Day 3 per SBI:** mcDNA extraction is performed with a recommended Maxiprep kit (Maxiprep handling takes about half a day, compared to Midiprep which can be done in an hour). SBI instructions recommend doubling the volumes of the Maxiprep buffers due to the large bacterial pellet size, which slows down handling further. As documented in the next section, the mcDNA yield was approximately 100 $\mu$g after Maxiprep extraction from a pellet produced from 400 ml of bacterial culture, while typical standard plasmid extraction with Midiprep from 45 ml of bacterial culture yields 400–600 $\mu$g of plasmid, explaining the one to two orders of magnitude higher endotoxin load per $\mu$g of DNA with the MC-Easy™ process. In our experience, endotoxin levels ranged from 47 to 121 EU/$\mu$g mcDNA. Notably, endotoxins are not mentioned in the MC-Easy™ Kit manual; thus, when researchers face issues with transfected cell culture survival, they might suspect something is wrong with the kit product, but with endotoxin testing being uncommon, they find the kit use cumbersome and yielding mcDNA toxic to sensitive cells.
- Quality controls follow, where mcDNA is cut by restriction enzymes, and gel electrophoresis is used to assess mcDNA purity. If the mcDNA quality is 'good', it

can be tested in transfection. However, if genomic DNA, parental plasmid DNA, or both are present, additional steps to remove these contaminations are necessary. SBI states: "If the parental DNA contamination is more than 10% higher than the minicircle DNA yield, you must restart again" (**Failure Point 4**). In our MC-Easy™ Kit validation experiments, in addition to gel electrophoresis, we used more precise whole plasmid sequencing by Plasmidsaurus, revealing the presence of recombined empty bacterial backbone at 4.2 kbp, parental plasmid at 11 kbp, and 5–6% bacterial DNA, making further purification steps necessary.

- We performed the 12-hour SBI Minicircle-safe DNase overnight treatment, heat inactivation of DNase, mcDNA isopropanol precipitation, and mcDNA purification removed impurities, with 34.5 $\mu$g of mcDNA remaining from the initial 109 $\mu$g (31.6% remaining). This added one more day to the processing time, and every time full plasmid sequencing was required for definitive analysis, an additional day wait was necessary, even with the fast Plasmidsaurus one-day turnaround. However, endotoxin levels were still high, ranging from 14.2–45.8 EU/$\mu$g mcDNA.
- While further purification of mcDNA could be possible using for instance methods listed in Almeida et al. [4] (SBI never cautioned about possible high endotoxin levels), this would require special reagents, additional equipment and labor, ultimately leading to an even lower yield of mcDNA.

##### Preparation and quality analysis of the MC DNA produced with MC-Easy™ kit

For validation of minicircle production, we used the same minicircle design as for the ABE8e base editing in the main article. We excised the mcDNA region from the parental ABE8e plasmid using restriction enzymes and inserted it into the SBI MC-Easy™ kit bacterial backbone plasmid following the kit instructions [3]. The designs of the parental plasmid *p-SBkit-ABE8e-parental-plasmid.gbk* (11071 bp) and the intended minicircle *SBkit-ABE8e-MC.gbk* (6936 bp) are available for review in the Supplementary Data folder *SB-Kit-Validation-Designs*. We followed the instructions in the MC-Easy™ kit and the manual [3] to produce the mcDNA. All operations were performed in duplicate (samples MC-1 and MC-2). At the end of the operations, we performed gel electrophoresis on all stages of the process, presented in Supplementary Figure **S11**. To aid in the identification of bands in mcDNAs, lanes L1–L3 show the intact and cut by restriction enzyme (*Fsp*I) SBkit-ABE8e-parental-plasmid and intact MC-Easy™ empty bacterial backbone plasmid. The further figures with whole plasmid sequencing provide much more precise information; observations from gel electrophoresis images and sequencing will be discussed further below.

The sequencing results in Supplementary Figure **S12**, produced by Plasmidsaurus Inc whole plasmid sequencing, show better quantitative analysis for the samples MC-1 and MC-2, compared to the gel image. Here we see that both samples contain relatively minor contamination with the 4.2 kbp MC-Easy™ recombined empty bacterial backbone, 11 kbp parental plasmid, and 5-6% *E. coli* DNA. This can also be seen in Supplementary Figure **S11** lanes L6–L7, showing weak bands matching the parental plasmid and the bacterial backbone plasmid, as well as a weak but noticeable diffuse glow. The figure shows high Nanodrop DNA purity for both samples; however, the yields are low, considering 400 ml of bacterial culture, 100 $\mu$g for MC-1 and 109 $\mu$g for MC-2.

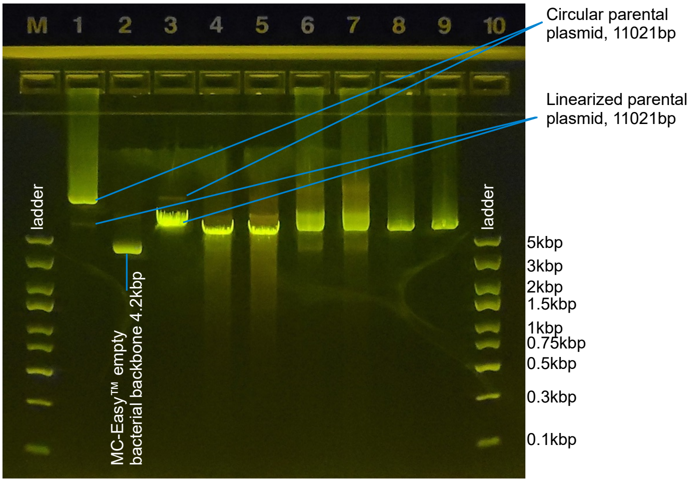


**Supplementary Figure S11**: **Gel electrophoresis of the MC DNA product produced with the MC-Easy™ kit** Lanes LM and L10 - ladder. L1 - parental plasmid; L2 - MC-Easy™ empty bacterial backbone plasmid; L3 - parental plasmid cut with restriction enzyme *Age*I, present only in MC but not in the bacterial backbone; L4 and L5 - the MC physical samples MC-1 and MC-2 cut with restriction enzyme *Age*I. L6 and L7 display the same MC-1 and MC-2 without restriction enzyme cutting, showing a clear picture; we can still see weak bands of parental plasmid and MC-Easy™ kit empty bacterial backbone. L8 and L9 are samples MC-1 and MC-2 after digestion with *Fsp*I restriction enzyme, present only in the bacterial backbone, followed by DNase digestion and column purification, finally showing the pure single MC band.

This compares poorly to the typical plasmid preparation using QIAGEN Midiprep Plus, which yields 400–600 $\mu$g of plasmid from 45 ml *E. coli* culture, as exemplified in Supplementary Figure **S20**.

Following the kit instructions [3], we used the restriction enzyme *Fsp*I, present only in the bacterial backbone but not in our intended mcDNA, and the minicircle-safe DNase provided by the MC-Easy$^{TM}$ kit to digest all non-circular or nicked DNA. This digestion stage was performed for 8 hours at 37°C, followed by isopropanol precipitation. The outcomes of this step are presented in Supplementary Figure **S13**, where we can see that the MC-Easy$^{TM}$ recombined empty bacterial backbone, parental plasmid, and *E. coli* DNA have been eliminated. This can also be seen in Supplementary Figure **S11** lanes L8–L9. This step resulted in high DNA purity; however, DNA loss was excessive, with only 15.2% of MC-1 and 31.6% of MC-2 remaining.

**Bacterial endotoxin levels of the mcDNA produced with MC-Easy™ kit**

While the MC-Easy$^{TM}$ kit manual [3] never mentions the word 'endotoxin', we hypothesized that the cell culture extinction we observed previously while transfecting using mcDNA produced with the MC-Easy$^{TM}$ kit might have been caused by high levels of

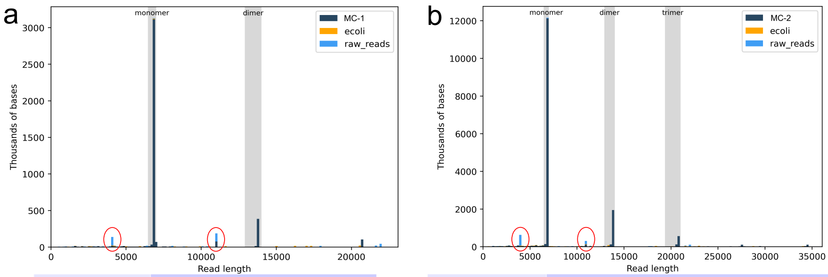


| | |
|---|---|
| MC-1 Plasmidsaurus sequencing 6936bp | 1-mer 2-mer 3-mer<br>moles 93.2 5.8 1.0<br>mass 86.5 10.8 2.8<br>**E. coli genomic contamination: 6.0%** |
| Yield and Nanodrop | Eluted in 200µl, 502ng/µl = 100µg DNA<br>Nanodrop: (260/280)=1.85;<br>(260/230) = 2.16 |

| | |
|---|---|
| MC-2 Plasmidsaurus sequencing 6936bp | 1-mer 2-mer 3-mer 4-mer 5-mer 6-8-mers 9-mer<br>moles 90.1 7.6 1.7 0.3 0.2 0.0 0.1<br>mass 79.7 13.4 4.5 1.1 0.9 0.0 0.4<br>**E. coli genomic contamination: 5.0%** |
| Yield and Nanodrop | Eluted in 200µl, 544ng/µl = 109µg DNA<br>Nanodrop: (260/280) = 1.83;<br>(260/230) = 2.16 |

**Supplementary Figure S12**: **MC DNA produced with MC-Easy™ kit, after Maxiprep isopropanol extraction.** (a) Sample MC-1; (b) sample MC-2. Both MCs perfectly match the sequence for the 6.9 kbp monomer with a minor fraction of multi-level concatemers. There is contamination with the 4.2 kbp MC-Easy™ recombined empty bacterial backbone plasmid and 11 kbp parental plasmid (circled in red), and 5–6% *E. coli* DNA.

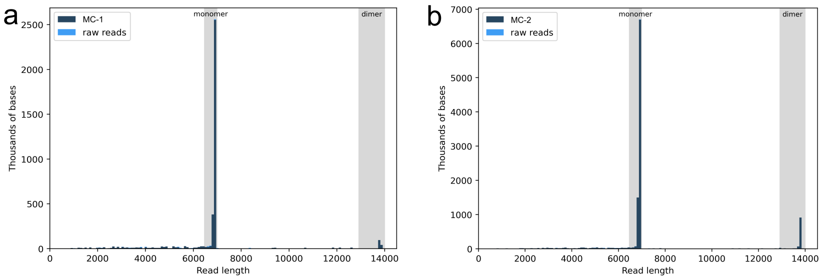


| | |
|---|---|
| MC-1 Plasmidsaurus sequencing 6936bp | (%) 1-mer 2-mer<br>moles 98.4 1.6<br>mass 96.8 3.2<br>**E. coli genomic contamination: 0.0%** |
| Yield and Nanodrop | 151.8ng/µl at 100µl = 15.2µg DNA<br>*(15.2% MC remaining after cleanup)*<br>Nanodrop: (260/280)=1.84;<br>(260/230) = 2.07 |

| | |
|---|---|
| MC-2 Plasmidsaurus sequencing 6936bp | (%) 1-mer 2-mer 3-mer<br>moles 93.7 5.9 0.4<br>mass 87.8 11.1 1.1<br>**E. coli genomic contamination: 0.0%** |
| Yield and Nanodrop | 344.7ng/µl at 100µl = 34.5µg DNA<br>*(31.6% MC remaining after cleanup)*<br>Nanodrop: (260/280) = 1.83;<br>(260/230) = 2.11 |

**Supplementary Figure S13**: **mcDNA produced with MC-Easy™ kit, after subsequent DNase digestion and cleanup.** (a) Sample MC-1; (b) sample MC-2. Both MCs again perfectly match the sequence for the 6.9 kbp monomer, and the fraction of concatemers has diminished. All initial DNA contaminations have been eliminated.

endotoxins. This hypothesis was based on the fact that the amount of cell membranes per unit of extracted circular DNA is significantly higher when 400 ml of bacterial culture is used to produce about 100 $\mu$g of DNA with MC-Easy versus 45 ml of bacterial culture to produce 400–600 $\mu$g of plasmid DNA with a typical Midiprep. To verify this, we used the Charles River Endosafe® nexgen-MCS endotoxin testing equipment and PTS2001F Limulus Amebocyte Lysate (LAL) endotoxin testing cartridges, certified for FDA validation of endotoxin levels, with the expertise of our affiliated Wyss Institute, which routinely performs such tests. Per the recommendation of Charles River technical support, we used a 10% dilution of Charles River BD100 dispersant, eliminating aggregation of endotoxins in DNA samples, which is critical for achieving consistent and reliable testing results. The PTS cartridges contain 4 lanes each for every individual sample, two for spike calibration and two for the endotoxin level measurement and cross-validation, resulting in highly accurate measurements. All tests were performed in triplicate.

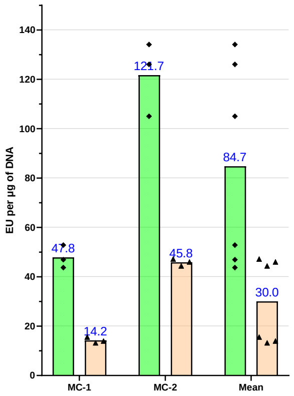


◆ Initial extraction and precipitation endotoxin levels

| | MC-Easy™ MC-1 EU/µg | MC-Easy™ MC-2 EU/µg |
|---|---|---|
| Phys Sample 1 | 43.7 | 134 |
| Phys Sample 2 | 52.8 | 126 |
| Phys Sample 3 | 46.9 | 105 |
| Mean | **47.9** | **121.7** |

▲ Digestion and cleanup endotoxin levels

| | MC-Easy™ MC-1 EU/µg | MC-Easy™ MC-2 EU/µg |
|---|---|---|
| Phys Sample 1 | 13.2 | 46.0 |
| Phys Sample 2 | 13.9 | 47.2 |
| Phys Sample 3 | 15.5 | 44.3 |
| Mean | **14.2** | **45.8** |

**Supplementary Figure S14**: **Endotoxin levels in mcDNA produced with MC-Easy™ kit.** The graphical representation of the endotoxin levels for samples MC-1 and MC-2. The tables on the right list the corresponding test values. See the corresponding test reports Supplementary Figure **S18**.

The results of this testing are presented in Supplementary Figure **S14**, which report critically elevated endotoxin levels. The endotoxin levels ranged from 47.9 to 121.7 EU/$\mu$g for the isopropanol-precipitated from Maxiprep mcDNA, with a mean value of 84.8 EU/$\mu$g. After digestion of the parental plasmid, empty bacterial backbone, and all other non-circular DNA, followed by cleanup, the endotoxin levels decreased by approximately threefold. However, the levels remained unacceptably high, ranging from 14.2 to 45.8 EU/$\mu$g, with a mean value of 30.0 EU/$\mu$g.

#### Conclusion

The System Biosciences mcDNA production kits are expensive, with costs of $1,072 for a 5-prep kit (Cat# MN920A-1) and $1,914 for a 10-prep kit (Cat# MN925A-1). Using ΦC31 pricing presented in the Discussion, producing a quality of mcDNA matching the MC-Easy™ Kit's 35 $\mu$g would require 125 $\mu$g of source plasmid and 125 $\mu$g of ΦC31 integrase, costing $10.24, rather than the cost of one preparation from the ten-pack MC-Easy™ Kit at $191.4. With the costs of remaining reagents, Maxiprep vs. Midiprep, and QIAGEN spin columns being approximately equivalent, this makes the Plasmid2MC method faster, less expensive, and more robust than using MC-Easy™ Kits.

The multi-step and multi-day preparation protocol lists numerous potential failure points where the manufacturer recommends restarting the production process if a parameter exceeds a specified limit [3]. Despite the relatively low yield of mcDNA, Maxiprep isopropanol extraction from bacterial culture contained traces of the parental plasmid, recombined empty parental bacterial backbone, *E. coli* genomic DNA, and high levels of endotoxins at 47–121 EU/$\mu$g (the MC-Easy$^{TM}$ kit manual and protocol do not mention endotoxins). Although the optional steps of DNase digestion and subsequent cleanup, as recommended by the manufacturer, removed DNA contamination at the cost of losing a fraction of mcDNA, endotoxin levels were only reduced by about threefold, remaining elevated at 14–45 EU/$\mu$g.

### The collection of the Charles River nexgen-MCS endotoxin test reports

This is the compendium of the endotoxin reports, referenced earlier in this Supplementary.

charles river

**Detailed LAL Endotoxin Report**

| | | | |
|---|---|---|---|
| **Serial Number/Bay:** MCS 24021012/1 | | **Start/End Temp (°C):** 37.0:37.0 | |
| **Sample Name:** ABE8e Plasmid | | | |
| **Sample ID:** N/A | | **Sample Lot:** N/A | **Dil./Conc.:** 2:1 mL/mL |
| **Dilution:** N/A | | | |
| **Sample Comments:** N/A | | | |
| **Cartridge Lot Number:** 4543158 | | **Cartridge Range:** 5-0.05 EU/mL | |
| **Calibration Code:** 515137484179 | | **Archived Spike Concentration:** 0.61 EU/mL | |
| **Range:** 151-774 seconds | | **Y-Intercept:** +2.427 | **Slope:** -0.355 |
| **Endotoxin Value:** 1.88 EU/mL | | **Sample CV Limit:** <25% | **Status:** VALID |
| **Endotoxin Limit:** No Limit | **Status:** N/A | **Spike CV Limit:** <25% | **Status:** VALID |
| **Alert Limit:** N/A | | **Spike Recovery Range:** 50-200% | **Status:** INVALID |
| **Cartridge Type:** LAL Cartridge | | **Performed With:** N/A | |

| SAMPLE DATA | | | | SPIKE DATA | | | | |
|---|---|---|---|---|---|---|---|---|
| Channel | Reaction Time | CV% | Sample Value | Channel | Reaction Time | CV% | Spike Value | Spike Recovery % |
| 1 | 162 | 4.2% | 1.88 EU/mL | 2 | <151 | 0.0% | >1.24 EU/mL | >203% |
| 3 | 172 | | | 4 | <151 | | | |

| | | | |
|---|---|---|---|
| **Serial Number/Bay:** MCS 24021013/2 | | **Start/End Temp (°C):** 37.0:37.0 | |
| **Sample Name:** ABE8e MC | | | |
| **Sample ID:** N/A | | **Sample Lot:** N/A | **Dil./Conc.:** 2:1 mL/mL |
| **Dilution:** N/A | | | |
| **Sample Comments:** N/A | | | |
| **Cartridge Lot Number:** 4543158 | | **Cartridge Range:** 5-0.05 EU/mL | |
| **Calibration Code:** 515137484179 | | **Archived Spike Concentration:** 0.61 EU/mL | |
| **Range:** 151-774 seconds | | **Y-Intercept:** +2.427 | **Slope:** -0.355 |
| **Endotoxin Value:** 0.029 EU/mL | | **Sample CV Limit:** <25% | **Status:** VALID |
| **Endotoxin Limit:** No Limit | **Status:** N/A | **Spike CV Limit:** <25% | **Status:** VALID |
| **Alert Limit:** N/A | | **Spike Recovery Range:** 50-200% | **Status:** VALID |
| **Cartridge Type:** LAL Cartridge | | **Performed With:** N/A | |

| SAMPLE DATA | | | | SPIKE DATA | | | | |
|---|---|---|---|---|---|---|---|---|
| Channel | Reaction Time | CV% | Sample Value | Channel | Reaction Time | CV% | Spike Value | Spike Recovery % |
| 1 | 744 | 1.5% | 0.029 EU/mL | 2 | 276 | 1.0% | 0.875 EU/mL | 143% |
| 3 | 728 | | | 4 | 272 | | | |

**Supplementary Figure S15**: **Preliminary test of Plasmid2MC endotoxin levels using PTS2005F cartridge.** Cartridge endotoxin range 5–0.05EU/ml, 2:1 dilution used.

charles river

**Detailed LAL Endotoxin Report**

**Source plasmid baseline endotoxin levels**

| | | | |
|---|---|---|---|
| **Serial Number/Bay:** MCS 24021012/1 | | **Start/End Temp (°C):** 37.0:37.0 | |
| **Sample Name:** ABE8e Plasmid Sample 1 | | | |
| **Sample ID:** N/A | | **Sample Lot:** N/A | **Dil./Conc.:** 1:5 mL/mL |
| **Dilution:** N/A | | | |
| **Sample Comments:** N/A | | | |
| **Cartridge Lot Number:** 3800176 | | **Cartridge Range:** 1-0.01 EU/mL | |
| **Calibration Code:** 116749143921 | | **Archived Spike Concentration:** 0.118 EU/mL | |
| **Range:** 167-891 seconds | | **Y-Intercept:** +2.223 | **Slope:** -0.364 |
| **Endotoxin Value:** 1.45 EU/mL | | **Sample CV Limit:** <25% | **Status:** VALID |
| **Endotoxin Limit:** No Limit | **Status:** N/A | **Spike CV Limit:** <25% | **Status:** VALID |
| **Alert Limit:** N/A | | **Spike Recovery Range:** 50-200% | **Status:** VALID |
| **Cartridge Type:** LAL Cartridge | | **Performed With:** N/A | |

| SAMPLE DATA | | | | SPIKE DATA | | | | |
|---|---|---|---|---|---|---|---|---|
| Channel | Reaction Time | CV% | Sample Value | Channel | Reaction Time | CV% | Spike Value | Spike Recovery % |
| 1 | 260 | 1.1% | 1.45 EU/mL | 2 | 222 | 3.1% | 0.140 EU/mL | 119% |
| 3 | 264 | | | 4 | 232 | | | |

| | | | |
|---|---|---|---|
| **Serial Number/Bay:** MCS 24021014/3 | | **Start/End Temp (°C):** 37.0:37.0 | |
| **Sample Name:** ABE8e Plasmid Sample 2 | | | |
| **Sample ID:** N/A | | **Sample Lot:** N/A | **Dil./Conc.:** 1:5 mL/mL |
| **Dilution:** N/A | | | |
| **Sample Comments:** N/A | | | |
| **Cartridge Lot Number:** 3800176 | | **Cartridge Range:** 1-0.01 EU/mL | |
| **Calibration Code:** 116749143921 | | **Archived Spike Concentration:** 0.118 EU/mL | |
| **Range:** 167-891 seconds | | **Y-Intercept:** +2.223 | **Slope:** -0.364 |
| **Endotoxin Value:** 1.58 EU/mL | | **Sample CV Limit:** <25% | **Status:** VALID |
| **Endotoxin Limit:** No Limit | **Status:** N/A | **Spike CV Limit:** <25% | **Status:** VALID |
| **Alert Limit:** N/A | | **Spike Recovery Range:** 50-200% | **Status:** VALID |
| **Cartridge Type:** LAL Cartridge | | **Performed With:** N/A | |

| SAMPLE DATA | | | | SPIKE DATA | | | | |
|---|---|---|---|---|---|---|---|---|
| Channel | Reaction Time | CV% | Sample Value | Channel | Reaction Time | CV% | Spike Value | Spike Recovery % |
| 1 | 278 | 13.4% | 1.58 EU/mL | 2 | 232 | 0.6% | 0.094 EU/mL | 80% |
| 3 | 230 | | | 4 | 230 | | | |

| | | | |
|---|---|---|---|
| **Serial Number/Bay:** MCS 24021012/1 | | **Start/End Temp (°C):** 37.0:37.0 | |
| **Sample Name:** ABE8e Plasmid Sample 3 | | | |
| **Sample ID:** N/A | | **Sample Lot:** N/A | **Dil./Conc.:** 1:5 mL/mL |
| **Dilution:** N/A | | | |
| **Sample Comments:** N/A | | | |
| **Cartridge Lot Number:** 3800176 | | **Cartridge Range:** 1-0.01 EU/mL | |
| **Calibration Code:** 116749143921 | | **Archived Spike Concentration:** 0.118 EU/mL | |
| **Range:** 167-891 seconds | | **Y-Intercept:** +2.223 | **Slope:** -0.364 |
| **Endotoxin Value:** 1.36 EU/mL | | **Sample CV Limit:** <25% | **Status:** VALID |
| **Endotoxin Limit:** No Limit | **Status:** N/A | **Spike CV Limit:** <25% | **Status:** VALID |
| **Alert Limit:** N/A | | **Spike Recovery Range:** 50-200% | **Status:** VALID |
| **Cartridge Type:** LAL Cartridge | | **Performed With:** N/A | |

| SAMPLE DATA | | | | SPIKE DATA | | | | |
|---|---|---|---|---|---|---|---|---|
| Channel | Reaction Time | CV% | Sample Value | Channel | Reaction Time | CV% | Spike Value | Spike Recovery % |
| 1 | 274 | 3.2% | 1.36 EU/mL | 2 | 234 | 6.3% | 0.174 EU/mL | 147% |
| 3 | 262 | | | 4 | 214 | | | |

**Endotoxin levels of mcDNA produced with Plasmid2MC method**

| | | | |
|---|---|---|---|
| **Serial Number/Bay:** MCS 24021014/3 | | **Start/End Temp (°C):** 37.0:37.0 | |
| **Sample Name:** ABE8e MC PhysSample 1 | | | |
| **Sample ID:** N/A | | **Sample Lot:** N/A | **Dil./Conc.:** 1:1 mL/mL |
| **Dilution:** N/A | | | |
| **Sample Comments:** N/A | | | |
| **Cartridge Lot Number:** 3800176 | | **Cartridge Range:** 1-0.01 EU/mL | |
| **Calibration Code:** 116749143921 | | **Archived Spike Concentration:** 0.118 EU/mL | |
| **Range:** 167-891 seconds | | **Y-Intercept:** +2.223 | **Slope:** -0.364 |
| **Endotoxin Value:** 0.0444 EU/mL | | **Sample CV Limit:** <25% | **Status:** VALID |
| **Endotoxin Limit:** No Limit | **Status:** N/A | **Spike CV Limit:** <25% | **Status:** VALID |
| **Alert Limit:** N/A | | **Spike Recovery Range:** 50-200% | **Status:** VALID |
| **Cartridge Type:** LAL Cartridge | | **Performed With:** N/A | |

| SAMPLE DATA | | | | SPIKE DATA | | | | |
|---|---|---|---|---|---|---|---|---|
| Channel | Reaction Time | CV% | Sample Value | Channel | Reaction Time | CV% | Spike Value | Spike Recovery % |
| 1 | 542 | 6.6% | 0.0444 EU/mL | 2 | 334 | 0.8% | 0.102 EU/mL | 86% |
| 3 | 494 | | | 4 | 338 | | | |

| | | | |
|---|---|---|---|
| **Serial Number/Bay:** MCS 24021015/4 | | **Start/End Temp (°C):** 37.0:37.0 | |
| **Sample Name:** ABE8e MC PhysSample 2 | | | |
| **Sample ID:** N/A | | **Sample Lot:** N/A | **Dil./Conc.:** 1:1 mL/mL |
| **Dilution:** N/A | | | |
| **Sample Comments:** N/A | | | |
| **Cartridge Lot Number:** 3800176 | | **Cartridge Range:** 1-0.01 EU/mL | |
| **Calibration Code:** 116749143921 | | **Archived Spike Concentration:** 0.118 EU/mL | |
| **Range:** 167-891 seconds | | **Y-Intercept:** +2.223 | **Slope:** -0.364 |
| **Endotoxin Value:** 0.0482 EU/mL | | **Sample CV Limit:** <25% | **Status:** VALID |
| **Endotoxin Limit:** No Limit | **Status:** N/A | **Spike CV Limit:** <25% | **Status:** VALID |
| **Alert Limit:** N/A | | **Spike Recovery Range:** 50-200% | **Status:** VALID |
| **Cartridge Type:** LAL Cartridge | | **Performed With:** N/A | |

| SAMPLE DATA | | | | SPIKE DATA | | | | |
|---|---|---|---|---|---|---|---|---|
| Channel | Reaction Time | CV% | Sample Value | Channel | Reaction Time | CV% | Spike Value | Spike Recovery % |
| 1 | 510 | 2.0% | 0.0482 EU/mL | 2 | 324 | 4.2% | 0.100 EU/mL | 85% |
| 3 | 496 | | | 4 | 344 | | | |

| | | | |
|---|---|---|---|
| **Serial Number/Bay:** MCS 24021014/3 | | **Start/End Temp (°C):** 37.0:37.0 | |
| **Sample Name:** ABE8e MC PhysSample 3 | | | |
| **Sample ID:** N/A | | **Sample Lot:** N/A | **Dil./Conc.:** 1:1 mL/mL |
| **Dilution:** N/A | | | |
| **Sample Comments:** N/A | | | |
| **Cartridge Lot Number:** 3800176 | | **Cartridge Range:** 1-0.01 EU/mL | |
| **Calibration Code:** 116749143921 | | **Archived Spike Concentration:** 0.118 EU/mL | |
| **Range:** 167-891 seconds | | **Y-Intercept:** +2.223 | **Slope:** -0.364 |
| **Endotoxin Value:** 0.0342 EU/mL | | **Sample CV Limit:** <25% | **Status:** VALID |
| **Endotoxin Limit:** No Limit | **Status:** N/A | **Spike CV Limit:** <25% | **Status:** VALID |
| **Alert Limit:** N/A | | **Spike Recovery Range:** 50-200% | **Status:** VALID |
| **Cartridge Type:** LAL Cartridge | | **Performed With:** N/A | |

| SAMPLE DATA | | | | SPIKE DATA | | | | |
|---|---|---|---|---|---|---|---|---|
| Channel | Reaction Time | CV% | Sample Value | Channel | Reaction Time | CV% | Spike Value | Spike Recovery % |
| 1 | 592 | 5.5% | 0.0342 EU/mL | 2 | 332 | 2.6% | 0.125 EU/mL | 106% |
| 3 | 548 | | | 4 | 320 | | | |

**Supplementary Figure S16**: **Endotoxin levels in mcDNA produced with Plasmid2MC method.** Top row: endotoxin levels in the source plasmid produced using QIAGEN MidPrep Plus; bottom row: endotoxin levels of the mcDNA produced from the above plasmids using Plasmid2MC method.

charles river

**Detailed LAL Endotoxin Report**

##### Source plasmid not containing SspI restriction enzyme recognition site: baseline endotoxin levels

| | | | |
|---|---|---|---|
| Serial Number/Bay: MCS 24021013/2 | | Start/End Temp (°C): 37.0:37.0 | |
| Sample Name: Plasmid for Digestion Test Sample 1 | | | |
| Sample ID: N/A | | Sample Lot: N/A | Dil./Conc.: 1:5 mL/mL |
| Dilution: N/A | | | |
| Sample Comments: N/A | | | |
| Cartridge Lot Number: 3771153 | | Cartridge Range: 1-0.01 EU/mL | |
| Calibration Code: 117556154382 | | Archived Spike Concentration: 0.126 EU/mL | |
| Range: 175-961 seconds | | Y-Intercept: +2.243 | Slope: -0.370 |
| Endotoxin Value: 0.133 EU/mL | | Sample CV Limit: <25% | Status: VALID |
| Endotoxin Limit: No Limit | Status: N/A | Spike CV Limit: <25% | Status: VALID |
| Alert Limit: N/A | | Spike Recovery Range: 50-200% | Status: VALID |
| Cartridge Type: LAL Cartridge | | Performed With: N/A | |

| SAMPLE DATA | | | | SPIKE DATA | | | | |
|---|---|---|---|---|---|---|---|---|
| Channel | Reaction Time | CV% | Sample Value | Channel | Reaction Time | CV% | Spike Value | Spike Recovery % |
| 1 | 674 | 0.8% | 0.133 EU/mL | 2 | 344 | 0.4% | 0.136 EU/mL | 108% |
| 3 | 666 | | | 4 | 342 | | | |

| | | | |
|---|---|---|---|
| Serial Number/Bay: MCS 24021012/1 | | Start/End Temp (°C): 37.0:37.0 | |
| Sample Name: Plasmid for Digestion Test Sample 2 | | | |
| Sample ID: N/A | | Sample Lot: N/A | Dil./Conc.: 1:1 mL/mL |
| Dilution: N/A | | | |
| Sample Comments: N/A | | | |
| Cartridge Lot Number: 3771153 | | Cartridge Range: 1-0.01 EU/mL | |
| Calibration Code: 117556154382 | | Archived Spike Concentration: 0.126 EU/mL | |
| Range: 175-961 seconds | | Y-Intercept: +2.243 | Slope: -0.370 |
| Endotoxin Value: 0.151 EU/mL | | Sample CV Limit: <25% | Status: VALID |
| Endotoxin Limit: No Limit | Status: N/A | Spike CV Limit: <25% | Status: VALID |
| Alert Limit: N/A | | Spike Recovery Range: 50-200% | Status: VALID |
| Cartridge Type: LAL Cartridge | | Performed With: N/A | |

| SAMPLE DATA | | | | SPIKE DATA | | | | |
|---|---|---|---|---|---|---|---|---|
| Channel | Reaction Time | CV% | Sample Value | Channel | Reaction Time | CV% | Spike Value | Spike Recovery % |
| 1 | 350 | 0.8% | 0.151 EU/mL | 2 | 300 | 2.3% | 0.072 EU/mL | 57% |
| 3 | 354 | | | 4 | 310 | | | |

| | | | |
|---|---|---|---|
| Serial Number/Bay: MCS 24021012/1 | | Start/End Temp (°C): 37.0:37.0 | |
| Sample Name: Plasmid for Digestion Test Sample 3 | | | |
| Sample ID: N/A | | Sample Lot: N/A | Dil./Conc.: 1:1 mL/mL |
| Dilution: N/A | | | |
| Sample Comments: N/A | | | |
| Cartridge Lot Number: 3771153 | | Cartridge Range: 1-0.01 EU/mL | |
| Calibration Code: 117556154382 | | Archived Spike Concentration: 0.126 EU/mL | |
| Range: 175-961 seconds | | Y-Intercept: +2.243 | Slope: -0.370 |
| Endotoxin Value: 0.227 EU/mL | | Sample CV Limit: <25% | Status: VALID |
| Endotoxin Limit: No Limit | Status: N/A | Spike CV Limit: <25% | Status: VALID |
| Alert Limit: N/A | | Spike Recovery Range: 50-200% | Status: VALID |
| Cartridge Type: LAL Cartridge | | Performed With: N/A | |

| SAMPLE DATA | | | | SPIKE DATA | | | | |
|---|---|---|---|---|---|---|---|---|
| Channel | Reaction Time | CV% | Sample Value | Channel | Reaction Time | CV% | Spike Value | Spike Recovery % |
| 1 | 302 | 0.5% | 0.227 EU/mL | 2 | 246 | 1.2% | 0.180 EU/mL | 143% |
| 3 | 304 | | | 4 | 242 | | | |

##### Digestion test with T5 Exonuclease and SspI restriction enzyme

| | | | |
|---|---|---|---|
| Serial Number/Bay: MCS 24021014/3 | | Start/End Temp (°C): 37.0:37.0 | |
| Sample Name: Plasmid Digestion Test T5+SspI PhysSample 1 | | | |
| Sample ID: N/A | | Sample Lot: N/A | Dil./Conc.: 1:1 mL/mL |
| Dilution: N/A | | | |
| Sample Comments: N/A | | | |
| Cartridge Lot Number: 3800176 | | Cartridge Range: 1-0.01 EU/mL | |
| Calibration Code: 116749143921 | | Archived Spike Concentration: 0.118 EU/mL | |
| Range: 167-891 seconds | | Y-Intercept: +2.223 | Slope: -0.364 |
| Endotoxin Value: 0.0355 EU/mL | | Sample CV Limit: <25% | Status: VALID |
| Endotoxin Limit: No Limit | Status: N/A | Spike CV Limit: <25% | Status: VALID |
| Alert Limit: N/A | | Spike Recovery Range: 50-200% | Status: VALID |
| Cartridge Type: LAL Cartridge | | Performed With: N/A | |

| SAMPLE DATA | | | | SPIKE DATA | | | | |
|---|---|---|---|---|---|---|---|---|
| Channel | Reaction Time | CV% | Sample Value | Channel | Reaction Time | CV% | Spike Value | Spike Recovery % |
| 1 | 558 | 1.0% | 0.0355 EU/mL | 2 | 308 | 13.3% | 0.106 EU/mL | 90% |
| 3 | 566 | | | 4 | 372 | | | |

| | | | |
|---|---|---|---|
| Serial Number/Bay: MCS 24021012/1 | | Start/End Temp (°C): 37.0:37.0 | |
| Sample Name: Plasmid Digestion Test T5+SspI PhysSample 2 | | | |
| Sample ID: N/A | | Sample Lot: N/A | Dil./Conc.: 1:1 mL/mL |
| Dilution: N/A | | | |
| Sample Comments: N/A | | | |
| Cartridge Lot Number: 3800176 | | Cartridge Range: 1-0.01 EU/mL | |
| Calibration Code: 116749143921 | | Archived Spike Concentration: 0.118 EU/mL | |
| Range: 167-891 seconds | | Y-Intercept: +2.223 | Slope: -0.364 |
| Endotoxin Value: 0.0348 EU/mL | | Sample CV Limit: <25% | Status: VALID |
| Endotoxin Limit: No Limit | Status: N/A | Spike CV Limit: <25% | Status: VALID |
| Alert Limit: N/A | | Spike Recovery Range: 50-200% | Status: VALID |
| Cartridge Type: LAL Cartridge | | Performed With: N/A | |

| SAMPLE DATA | | | | SPIKE DATA | | | | |
|---|---|---|---|---|---|---|---|---|
| Channel | Reaction Time | CV% | Sample Value | Channel | Reaction Time | CV% | Spike Value | Spike Recovery % |
| 1 | 580 | 3.5% | 0.0348 EU/mL | 2 | 342 | 2.0% | 0.099 EU/mL | 84% |
| 3 | 552 | | | 4 | 352 | | | |

| | | | |
|---|---|---|---|
| Serial Number/Bay: MCS 24021012/1 | | Start/End Temp (°C): 37.0:37.0 | |
| Sample Name: Plasmid Digestion Test T5+SspI PhysSample 3 | | | |
| Sample ID: N/A | | Sample Lot: N/A | Dil./Conc.: 1:1 mL/mL |
| Dilution: N/A | | | |
| Sample Comments: N/A | | | |
| Cartridge Lot Number: 3800176 | | Cartridge Range: 1-0.01 EU/mL | |
| Calibration Code: 116749143921 | | Archived Spike Concentration: 0.118 EU/mL | |
| Range: 167-891 seconds | | Y-Intercept: +2.223 | Slope: -0.364 |
| Endotoxin Value: 0.0466 EU/mL | | Sample CV Limit: <25% | Status: VALID |
| Endotoxin Limit: No Limit | Status: N/A | Spike CV Limit: <25% | Status: VALID |
| Alert Limit: N/A | | Spike Recovery Range: 50-200% | Status: VALID |
| Cartridge Type: LAL Cartridge | | Performed With: N/A | |

| SAMPLE DATA | | | | SPIKE DATA | | | | |
|---|---|---|---|---|---|---|---|---|
| Channel | Reaction Time | CV% | Sample Value | Channel | Reaction Time | CV% | Spike Value | Spike Recovery % |
| 1 | 518 | 2.5% | 0.0466 EU/mL | 2 | 330 | 5.8% | 0.125 EU/mL | 106% |
| 3 | 500 | | | 4 | 304 | | | |

##### Digestion test with T5 Exonuclease and SspI restriction enzyme, identical to above, followed by Proteinase K protein digestion

| | | | |
|---|---|---|---|
| Serial Number/Bay: MCS 24021013/2 | | Start/End Temp (°C): 37.0:37.0 | |
| Sample Name: Plasmid Digestion Test T5+SspI followed by ProteinaseK PhysSample 1 | | | |
| Sample ID: N/A | | Sample Lot: N/A | Dil./Conc.: 1:1 mL/mL |
| Dilution: N/A | | | |
| Sample Comments: N/A | | | |
| Cartridge Lot Number: 3800176 | | Cartridge Range: 1-0.01 EU/mL | |
| Calibration Code: 116749143921 | | Archived Spike Concentration: 0.118 EU/mL | |
| Range: 167-891 seconds | | Y-Intercept: +2.223 | Slope: -0.364 |
| Endotoxin Value: 0.158 EU/mL | | Sample CV Limit: <25% | Status: VALID |
| Endotoxin Limit: No Limit | Status: N/A | Spike CV Limit: <25% | Status: VALID |
| Alert Limit: N/A | | Spike Recovery Range: 50-200% | Status: VALID |
| Cartridge Type: LAL Cartridge | | Performed With: N/A | |

| SAMPLE DATA | | | | SPIKE DATA | | | | |
|---|---|---|---|---|---|---|---|---|
| Channel | Reaction Time | CV% | Sample Value | Channel | Reaction Time | CV% | Spike Value | Spike Recovery % |
| 1 | 332 | 2.2% | 0.158 EU/mL | 2 | 278 | 4.2% | 0.109 EU/mL | 93% |
| 3 | 322 | | | 4 | 262 | | | |

| | | | |
|---|---|---|---|
| Serial Number/Bay: MCS 24021013/2 | | Start/End Temp (°C): 37.0:37.0 | |
| Sample Name: Plasmid Digestion Test T5+SspI followed by ProteinaseK PhysSample 2 | | | |
| Sample ID: N/A | | Sample Lot: N/A | Dil./Conc.: 1:1 mL/mL |
| Dilution: N/A | | | |
| Sample Comments: N/A | | | |
| Cartridge Lot Number: 3800176 | | Cartridge Range: 1-0.01 EU/mL | |
| Calibration Code: 116749143921 | | Archived Spike Concentration: 0.118 EU/mL | |
| Range: 167-891 seconds | | Y-Intercept: +2.223 | Slope: -0.364 |
| Endotoxin Value: 0.169 EU/mL | | Sample CV Limit: <25% | Status: VALID |
| Endotoxin Limit: No Limit | Status: N/A | Spike CV Limit: <25% | Status: VALID |
| Alert Limit: N/A | | Spike Recovery Range: 50-200% | Status: VALID |
| Cartridge Type: LAL Cartridge | | Performed With: N/A | |

| SAMPLE DATA | | | | SPIKE DATA | | | | |
|---|---|---|---|---|---|---|---|---|
| Channel | Reaction Time | CV% | Sample Value | Channel | Reaction Time | CV% | Spike Value | Spike Recovery % |
| 1 | 326 | 3.1% | 0.169 EU/mL | 2 | 260 | 0.0% | 0.127 EU/mL | 108% |
| 3 | 312 | | | 4 | 260 | | | |

| | | | |
|---|---|---|---|
| Serial Number/Bay: MCS 24021013/2 | | Start/End Temp (°C): 37.0:37.0 | |
| Sample Name: Plasmid Digestion Test T5+SspI followed by ProteinaseK PhysSample 3 | | | |
| Sample ID: N/A | | Sample Lot: N/A | Dil./Conc.: 1:1 mL/mL |
| Dilution: N/A | | | |
| Sample Comments: N/A | | | |
| Cartridge Lot Number: 3800176 | | Cartridge Range: 1-0.01 EU/mL | |
| Calibration Code: 116749143921 | | Archived Spike Concentration: 0.118 EU/mL | |
| Range: 167-891 seconds | | Y-Intercept: +2.223 | Slope: -0.364 |
| Endotoxin Value: 0.136 EU/mL | | Sample CV Limit: <25% | Status: VALID |
| Endotoxin Limit: No Limit | Status: N/A | Spike CV Limit: <25% | Status: VALID |
| Alert Limit: N/A | | Spike Recovery Range: 50-200% | Status: VALID |
| Cartridge Type: LAL Cartridge | | Performed With: N/A | |

| SAMPLE DATA | | | | SPIKE DATA | | | | |
|---|---|---|---|---|---|---|---|---|
| Channel | Reaction Time | CV% | Sample Value | Channel | Reaction Time | CV% | Spike Value | Spike Recovery % |
| 1 | 358 | 5.3% | 0.136 EU/mL | 2 | 270 | 3.6% | 0.113 EU/mL | 96% |
| 3 | 332 | | | 4 | 284 | | | |

**Supplementary Figure S17**: **Reports used for the testing of endotoxin elimination during the digestion stage.** These tests were performed to validate the level reduction assisted by endotoxins being carried away by the restriction enzymes and endonuclease.

### Detailed LAL Endotoxin Report

#### Sample MC-1 endotoxin level measurements after Maxiprep isopropanol precipitation

| | | | |
|---|---|---|---|
| Serial Number/Bay: MCS 24021012/1 | | Start/End Temp (°C): 37.0:37.0 | |
| Sample Name: SB Kit MC PhysSample 1a | | | |
| Sample ID: N/A | | Sample Lot: N/A | Dil./Conc.: 1:500 mL/mL |
| Dilution: N/A | | | |
| Sample Comments: N/A | | | |
| Cartridge Lot Number: 3800176 | | Cartridge Range: 1-0.01 EU/mL | |
| Calibration Code: 116749143921 | | Archived Spike Concentration: 0.118 EU/mL | |
| Range: 167-891 seconds | | Y-Intercept: +2.223 | Slope: -0.364 |
| Endotoxin Value: 43.7 EU/mL | | Sample CV Limit: <25% | Status: VALID |
| Endotoxin Limit: No Limit | Status: N/A | Spike CV Limit: <25% | Status: VALID |
| Alert Limit: N/A | | Spike Recovery Range: 50-200% | Status: VALID |
| Cartridge Type: LAL Cartridge | | Performed With: N/A | |

| SAMPLE DATA | | | | SPIKE DATA | | | | |
|---|---|---|---|---|---|---|---|---|
| Channel | Reaction Time | CV% | Sample Value | Channel | Reaction Time | CV% | Spike Value | Spike Recovery % |
| 1 | 430 | 8.7% | 43.7 EU/mL | 2 | 316 | 1.8% | 0.080 EU/mL | 68% |
| 3 | 380 | | | 4 | 324 | | | |

| | | | |
|---|---|---|---|
| Serial Number/Bay: MCS 24021013/2 | | Start/End Temp (°C): 37.0:37.0 | |
| Sample Name: SB Kit MC PhysSample 1b | | | |
| Sample ID: N/A | | Sample Lot: N/A | Dil./Conc.: 1:200 mL/mL |
| Dilution: N/A | | | |
| Sample Comments: N/A | | | |
| Cartridge Lot Number: 3800176 | | Cartridge Range: 1-0.01 EU/mL | |
| Calibration Code: 116749143921 | | Archived Spike Concentration: 0.118 EU/mL | |
| Range: 167-891 seconds | | Y-Intercept: +2.223 | Slope: -0.364 |
| Endotoxin Value: 52.8 EU/mL | | Sample CV Limit: <25% | Status: VALID |
| Endotoxin Limit: No Limit | Status: N/A | Spike CV Limit: <25% | Status: VALID |
| Alert Limit: N/A | | Spike Recovery Range: 50-200% | Status: VALID |
| Cartridge Type: LAL Cartridge | | Performed With: N/A | |

| SAMPLE DATA | | | | SPIKE DATA | | | | |
|---|---|---|---|---|---|---|---|---|
| Channel | Reaction Time | CV% | Sample Value | Channel | Reaction Time | CV% | Spike Value | Spike Recovery % |
| 1 | 274 | 1.6% | 52.8 EU/mL | 2 | 236 | 2.9% | 0.101 EU/mL | 85% |
| 3 | 268 | | | 4 | 246 | | | |

| | | | |
|---|---|---|---|
| Serial Number/Bay: MCS 24021014/3 | | Start/End Temp (°C): 37.0:37.0 | |
| Sample Name: SB Kit MC PhysSample 1c | | | |
| Sample ID: N/A | | Sample Lot: N/A | Dil./Conc.: 1:200 mL/mL |
| Dilution: N/A | | | |
| Sample Comments: N/A | | | |
| Cartridge Lot Number: 3800176 | | Cartridge Range: 1-0.01 EU/mL | |
| Calibration Code: 116749143921 | | Archived Spike Concentration: 0.118 EU/mL | |
| Range: 167-891 seconds | | Y-Intercept: +2.223 | Slope: -0.364 |
| Endotoxin Value: 46.9 EU/mL | | Sample CV Limit: <25% | Status: VALID |
| Endotoxin Limit: No Limit | Status: N/A | Spike CV Limit: <25% | Status: VALID |
| Alert Limit: N/A | | Spike Recovery Range: 50-200% | Status: VALID |
| Cartridge Type: LAL Cartridge | | Performed With: N/A | |

| SAMPLE DATA | | | | SPIKE DATA | | | | |
|---|---|---|---|---|---|---|---|---|
| Channel | Reaction Time | CV% | Sample Value | Channel | Reaction Time | CV% | Spike Value | Spike Recovery % |
| 1 | 294 | 5.5% | 46.9 EU/mL | 2 | 238 | 3.5% | 0.118 EU/mL | 100% |
| 3 | 272 | | | 4 | 250 | | | |

#### Sample MC-2 endotoxin level measurements after Maxiprep isopropanol precipitation

| | | | |
|---|---|---|---|
| Serial Number/Bay: MCS 24021015/4 | | Start/End Temp (°C): 37.0:37.0 | |
| Sample Name: SB Kit MC PhysSample 2a | | | |
| Sample ID: N/A | | Sample Lot: N/A | Dil./Conc.: 1:500 mL/mL |
| Dilution: N/A | | | |
| Sample Comments: N/A | | | |
| Cartridge Lot Number: 3800176 | | Cartridge Range: 1-0.01 EU/mL | |
| Calibration Code: 116749143921 | | Archived Spike Concentration: 0.118 EU/mL | |
| Range: 167-891 seconds | | Y-Intercept: +2.223 | Slope: -0.364 |
| Endotoxin Value: 134 EU/mL | | Sample CV Limit: <25% | Status: VALID |
| Endotoxin Limit: No Limit | Status: N/A | Spike CV Limit: <25% | Status: VALID |
| Alert Limit: N/A | | Spike Recovery Range: 50-200% | Status: VALID |
| Cartridge Type: LAL Cartridge | | Performed With: N/A | |

| SAMPLE DATA | | | | SPIKE DATA | | | | |
|---|---|---|---|---|---|---|---|---|
| Channel | Reaction Time | CV% | Sample Value | Channel | Reaction Time | CV% | Spike Value | Spike Recovery % |
| 1 | 270 | 0.0% | 134 EU/mL | 2 | 222 | 5.5% | 0.143 EU/mL | 121% |
| 3 | 270 | | | 4 | 240 | | | |

| | | | |
|---|---|---|---|
| Serial Number/Bay: MCS 24021016/5 | | Start/End Temp (°C): 37.0:37.0 | |
| Sample Name: SB Kit MC PhysSample 2c | | | |
| Sample ID: N/A | | Sample Lot: N/A | Dil./Conc.: 1:500 mL/mL |
| Dilution: N/A | | | |
| Sample Comments: N/A | | | |
| Cartridge Lot Number: 3800176 | | Cartridge Range: 1-0.01 EU/mL | |
| Calibration Code: 116749143921 | | Archived Spike Concentration: 0.118 EU/mL | |
| Range: 167-891 seconds | | Y-Intercept: +2.223 | Slope: -0.364 |
| Endotoxin Value: 126 EU/mL | | Sample CV Limit: <25% | Status: VALID |
| Endotoxin Limit: No Limit | Status: N/A | Spike CV Limit: <25% | Status: VALID |
| Alert Limit: N/A | | Spike Recovery Range: 50-200% | Status: VALID |
| Cartridge Type: LAL Cartridge | | Performed With: N/A | |

| SAMPLE DATA | | | | SPIKE DATA | | | | |
|---|---|---|---|---|---|---|---|---|
| Channel | Reaction Time | CV% | Sample Value | Channel | Reaction Time | CV% | Spike Value | Spike Recovery % |
| 1 | 284 | 4.1% | 126 EU/mL | 2 | 242 | 2.4% | 0.126 EU/mL | 107% |
| 3 | 268 | | | 4 | 234 | | | |

| | | | |
|---|---|---|---|
| Serial Number/Bay: MCS 24021012/1 | | Start/End Temp (°C): 37.0:37.0 | |
| Sample Name: SB Kit MC PhysSample 2d | | | |
| Sample ID: N/A | | Sample Lot: N/A | Dil./Conc.: 1:500 mL/mL |
| Dilution: N/A | | | |
| Sample Comments: N/A | | | |
| Cartridge Lot Number: 3771153 | | Cartridge Range: 1-0.01 EU/mL | |
| Calibration Code: 117556154382 | | Archived Spike Concentration: 0.126 EU/mL | |
| Range: 175-961 seconds | | Y-Intercept: +2.243 | Slope: -0.370 |
| Endotoxin Value: 105 EU/mL | | Sample CV Limit: <25% | Status: VALID |
| Endotoxin Limit: No Limit | Status: N/A | Spike CV Limit: <25% | Status: VALID |
| Alert Limit: N/A | | Spike Recovery Range: 50-200% | Status: VALID |
| Cartridge Type: LAL Cartridge | | Performed With: N/A | |

| SAMPLE DATA | | | | SPIKE DATA | | | | |
|---|---|---|---|---|---|---|---|---|
| Channel | Reaction Time | CV% | Sample Value | Channel | Reaction Time | CV% | Spike Value | Spike Recovery % |
| 1 | 320 | 3.6% | 105 EU/mL | 2 | 252 | 1.1% | 0.172 EU/mL | 136% |
| 3 | 304 | | | 4 | 248 | | | |

#### Sample MC-1 endotoxin level measurements after optional cleanup steps

| | | | |
|---|---|---|---|
| Serial Number/Bay: MCS 24021015/4 | | Start/End Temp (°C): 37.0:37.0 | |
| Sample Name: SB Kit MC Cleaned PhysSample 1a | | | |
| Sample ID: N/A | | Sample Lot: N/A | Dil./Conc.: 1:500 mL/mL |
| Dilution: N/A | | | |
| Sample Comments: N/A | | | |
| Cartridge Lot Number: 3771153 | | Cartridge Range: 1-0.01 EU/mL | |
| Calibration Code: 117556154382 | | Archived Spike Concentration: 0.126 EU/mL | |
| Range: 175-961 seconds | | Y-Intercept: +2.243 | Slope: -0.370 |
| Endotoxin Value: 46.0 EU/mL | | Sample CV Limit: <25% | Status: VALID |
| Endotoxin Limit: No Limit | Status: N/A | Spike CV Limit: <25% | Status: VALID |
| Alert Limit: N/A | | Spike Recovery Range: 50-200% | Status: VALID |
| Cartridge Type: LAL Cartridge | | Performed With: N/A | |

| SAMPLE DATA | | | | SPIKE DATA | | | | |
|---|---|---|---|---|---|---|---|---|
| Channel | Reaction Time | CV% | Sample Value | Channel | Reaction Time | CV% | Spike Value | Spike Recovery % |
| 1 | 422 | 0.3% | 46.0 EU/mL | 2 | 308 | 6.7% | 0.154 EU/mL | 122% |
| 3 | 424 | | | 4 | 280 | | | |

| | | | |
|---|---|---|---|
| Serial Number/Bay: MCS 24021013/2 | | Start/End Temp (°C): 37.0:37.0 | |
| Sample Name: SB Kit MC Cleaned PhysSample 1b | | | |
| Sample ID: N/A | | Sample Lot: N/A | Dil./Conc.: 1:500 mL/mL |
| Dilution: N/A | | | |
| Sample Comments: N/A | | | |
| Cartridge Lot Number: 3771153 | | Cartridge Range: 1-0.01 EU/mL | |
| Calibration Code: 117556154382 | | Archived Spike Concentration: 0.126 EU/mL | |
| Range: 175-961 seconds | | Y-Intercept: +2.243 | Slope: -0.370 |
| Endotoxin Value: 47.2 EU/mL | | Sample CV Limit: <25% | Status: VALID |
| Endotoxin Limit: No Limit | Status: N/A | Spike CV Limit: <25% | Status: VALID |
| Alert Limit: N/A | | Spike Recovery Range: 50-200% | Status: VALID |
| Cartridge Type: LAL Cartridge | | Performed With: N/A | |

| SAMPLE DATA | | | | SPIKE DATA | | | | |
|---|---|---|---|---|---|---|---|---|
| Channel | Reaction Time | CV% | Sample Value | Channel | Reaction Time | CV% | Spike Value | Spike Recovery % |
| 1 | 420 | 0.3% | 47.2 EU/mL | 2 | 296 | 2.4% | 0.158 EU/mL | 126% |
| 3 | 418 | | | 4 | 286 | | | |

| | | | |
|---|---|---|---|
| Serial Number/Bay: MCS 24021014/3 | | Start/End Temp (°C): 37.0:37.0 | |
| Sample Name: SB Kit MC Cleaned PhysSample 1c | | | |
| Sample ID: N/A | | Sample Lot: N/A | Dil./Conc.: 1:500 mL/mL |
| Dilution: N/A | | | |
| Sample Comments: N/A | | | |
| Cartridge Lot Number: 3771153 | | Cartridge Range: 1-0.01 EU/mL | |
| Calibration Code: 117556154382 | | Archived Spike Concentration: 0.126 EU/mL | |
| Range: 175-961 seconds | | Y-Intercept: +2.243 | Slope: -0.370 |
| Endotoxin Value: 44.3 EU/mL | | Sample CV Limit: <25% | Status: VALID |
| Endotoxin Limit: No Limit | Status: N/A | Spike CV Limit: <25% | Status: VALID |
| Alert Limit: N/A | | Spike Recovery Range: 50-200% | Status: VALID |
| Cartridge Type: LAL Cartridge | | Performed With: N/A | |

| SAMPLE DATA | | | | SPIKE DATA | | | | |
|---|---|---|---|---|---|---|---|---|
| Channel | Reaction Time | CV% | Sample Value | Channel | Reaction Time | CV% | Spike Value | Spike Recovery % |
| 1 | 436 | 2.3% | 44.3 EU/mL | 2 | 296 | 2.4% | 0.164 EU/mL | 130% |
| 3 | 422 | | | 4 | 286 | | | |

#### Sample MC-2 endotoxin level measurements after optional cleanup steps

| | | | |
|---|---|---|---|
| Serial Number/Bay: MCS 24021013/2 | | Start/End Temp (°C): 37.0:37.0 | |
| Sample Name: SB Kit MC Cleaned PhysSample 2a | | | |
| Sample ID: N/A | | Sample Lot: N/A | Dil./Conc.: 1:500 mL/mL |
| Dilution: N/A | | | |
| Sample Comments: N/A | | | |
| Cartridge Lot Number: 3771153 | | Cartridge Range: 1-0.01 EU/mL | |
| Calibration Code: 117556154382 | | Archived Spike Concentration: 0.126 EU/mL | |
| Range: 175-961 seconds | | Y-Intercept: +2.243 | Slope: -0.370 |
| Endotoxin Value: 13.2 EU/mL | | Sample CV Limit: <25% | Status: VALID |
| Endotoxin Limit: No Limit | Status: N/A | Spike CV Limit: <25% | Status: VALID |
| Alert Limit: N/A | | Spike Recovery Range: 50-200% | Status: VALID |
| Cartridge Type: LAL Cartridge | | Performed With: N/A | |

| SAMPLE DATA | | | | SPIKE DATA | | | | |
|---|---|---|---|---|---|---|---|---|
| Channel | Reaction Time | CV% | Sample Value | Channel | Reaction Time | CV% | Spike Value | Spike Recovery % |
| 1 | 658 | 2.9% | 13.2 EU/mL | 2 | 378 | 1.9% | 0.103 EU/mL | 82% |
| 3 | 686 | | | 4 | 368 | | | |

| | | | |
|---|---|---|---|
| Serial Number/Bay: MCS 24021016/5 | | Start/End Temp (°C): 37.0:37.0 | |
| Sample Name: SB Kit MC Cleaned PhysSample 2b | | | |
| Sample ID: N/A | | Sample Lot: N/A | Dil./Conc.: 1:50 mL/mL |
| Dilution: N/A | | | |
| Sample Comments: N/A | | | |
| Cartridge Lot Number: 3771153 | | Cartridge Range: 1-0.01 EU/mL | |
| Calibration Code: 117556154382 | | Archived Spike Concentration: 0.126 EU/mL | |
| Range: 175-961 seconds | | Y-Intercept: +2.243 | Slope: -0.370 |
| Endotoxin Value: 13.9 EU/mL | | Sample CV Limit: <25% | Status: VALID |
| Endotoxin Limit: No Limit | Status: N/A | Spike CV Limit: <25% | Status: VALID |
| Alert Limit: N/A | | Spike Recovery Range: 50-200% | Status: VALID |
| Cartridge Type: LAL Cartridge | | Performed With: N/A | |

| SAMPLE DATA | | | | SPIKE DATA | | | | |
|---|---|---|---|---|---|---|---|---|
| Channel | Reaction Time | CV% | Sample Value | Channel | Reaction Time | CV% | Spike Value | Spike Recovery % |
| 1 | 276 | 2.5% | 13.9 EU/mL | 2 | 232 | 0.6% | 0.194 EU/mL | 154% |
| 3 | 286 | | | 4 | 230 | | | |

| | | | |
|---|---|---|---|
| Serial Number/Bay: MCS 24021015/4 | | Start/End Temp (°C): 37.0:37.0 | |
| Sample Name: SB Kit MC Cleaned PhysSample 2c | | | |
| Sample ID: N/A | | Sample Lot: N/A | Dil./Conc.: 1:50 mL/mL |
| Dilution: N/A | | | |
| Sample Comments: N/A | | | |
| Cartridge Lot Number: 3771153 | | Cartridge Range: 1-0.01 EU/mL | |
| Calibration Code: 117556154382 | | Archived Spike Concentration: 0.126 EU/mL | |
| Range: 175-961 seconds | | Y-Intercept: +2.243 | Slope: -0.370 |
| Endotoxin Value: 15.5 EU/mL | | Sample CV Limit: <25% | Status: VALID |
| Endotoxin Limit: No Limit | Status: N/A | Spike CV Limit: <25% | Status: VALID |
| Alert Limit: N/A | | Spike Recovery Range: 50-200% | Status: VALID |
| Cartridge Type: LAL Cartridge | | Performed With: N/A | |

| SAMPLE DATA | | | | SPIKE DATA | | | | |
|---|---|---|---|---|---|---|---|---|
| Channel | Reaction Time | CV% | Sample Value | Channel | Reaction Time | CV% | Spike Value | Spike Recovery % |
| 1 | 266 | 2.1% | 15.5 EU/mL | 2 | 230 | 3.8% | 0.203 EU/mL | 161% |
| 3 | 274 | | | 4 | 218 | | | |

**Supplementary Figure S18**: **Endotoxin levels in mcDNA produced with MC-Easy™ kit.** The graphical representation of the endotoxin levels for samples MC-1 and MC-2. The tables on the right list the corresponding test values.

charles river

**Detailed LAL Endotoxin Report**

**Endotoxin levels in plasmid sample 2 prepared using QIAGEN MidiPrep Plus kit**

| | | |
|---|---|---|
| **Serial Number/Bay:** MCS 24021015/4 | **Start/End Temp (°C):** 37.0:37.0 | |
| **Sample Name:** ABE8e Plasmid-2 Sample 1 | | |
| **Sample ID:** N/A | **Sample Lot:** N/A | **Dil./Conc.:** 1:5 mL/mL |
| **Dilution:** N/A | | |
| **Sample Comments:** N/A | | |
| **Cartridge Lot Number:** 3771153 | **Cartridge Range:** 1-0.01 EU/mL | |
| **Calibration Code:** 117556154382 | **Archived Spike Concentration:** 0.126 EU/mL | |
| **Range:** 175-961 seconds | **Y-Intercept:** +2.243 | **Slope:** -0.370 |
| **Endotoxin Value:** 0.628 EU/mL | **Sample CV Limit:** <25% | **Status:** VALID |
| **Endotoxin Limit:** No Limit **Status:** N/A | **Spike CV Limit:** <25% | **Status:** VALID |
| **Alert Limit:** N/A | **Spike Recovery Range:** 50-200% | **Status:** VALID |
| **Cartridge Type:** LAL Cartridge | **Performed With:** N/A | |

| SAMPLE DATA | | | | SPIKE DATA | | | | |
|---|---|---|---|---|---|---|---|---|
| Channel | Reaction Time | CV% | Sample Value | Channel | Reaction Time | CV% | Spike Value | Spike Recovery % |
| 1 | 386 | 3.4% | 0.628 EU/mL | 2 | 282 | 2.0% | 0.139 EU/mL | 111% |
| 3 | 368 | | | 4 | 290 | | | |

| | | |
|---|---|---|
| **Serial Number/Bay:** MCS 24021013/2 | **Start/End Temp (°C):** 37.0:37.0 | |
| **Sample Name:** ABE8e Plasmid-2 Sample 2 | | |
| **Sample ID:** N/A | **Sample Lot:** N/A | **Dil./Conc.:** 1:5 mL/mL |
| **Dilution:** N/A | | |
| **Sample Comments:** N/A | | |
| **Cartridge Lot Number:** 3771153 | **Cartridge Range:** 1-0.01 EU/mL | |
| **Calibration Code:** 117556154382 | **Archived Spike Concentration:** 0.126 EU/mL | |
| **Range:** 175-961 seconds | **Y-Intercept:** +2.243 | **Slope:** -0.370 |
| **Endotoxin Value:** 0.550 EU/mL | **Sample CV Limit:** <25% | **Status:** VALID |
| **Endotoxin Limit:** No Limit **Status:** N/A | **Spike CV Limit:** <25% | **Status:** VALID |
| **Alert Limit:** N/A | **Spike Recovery Range:** 50-200% | **Status:** VALID |
| **Cartridge Type:** LAL Cartridge | **Performed With:** N/A | |

| SAMPLE DATA | | | | SPIKE DATA | | | | |
|---|---|---|---|---|---|---|---|---|
| Channel | Reaction Time | CV% | Sample Value | Channel | Reaction Time | CV% | Spike Value | Spike Recovery % |
| 1 | 394 | 0.7% | 0.550 EU/mL | 2 | 278 | 2.0% | 0.165 EU/mL | 131% |
| 3 | 398 | | | 4 | 286 | | | |

| | | |
|---|---|---|
| **Serial Number/Bay:** MCS 24021014/3 | **Start/End Temp (°C):** 37.0:37.0 | |
| **Sample Name:** ABE8e Plasmid-2 Sample 3 | | |
| **Sample ID:** N/A | **Sample Lot:** N/A | **Dil./Conc.:** 1:5 mL/mL |
| **Dilution:** N/A | | |
| **Sample Comments:** N/A | | |
| **Cartridge Lot Number:** 3771153 | **Cartridge Range:** 1-0.01 EU/mL | |
| **Calibration Code:** 117556154382 | **Archived Spike Concentration:** 0.126 EU/mL | |
| **Range:** 175-961 seconds | **Y-Intercept:** +2.243 | **Slope:** -0.370 |
| **Endotoxin Value:** 0.528 EU/mL | **Sample CV Limit:** <25% | **Status:** VALID |
| **Endotoxin Limit:** No Limit **Status:** N/A | **Spike CV Limit:** <25% | **Status:** VALID |
| **Alert Limit:** N/A | **Spike Recovery Range:** 50-200% | **Status:** VALID |
| **Cartridge Type:** LAL Cartridge | **Performed With:** N/A | |

| SAMPLE DATA | | | | SPIKE DATA | | | | |
|---|---|---|---|---|---|---|---|---|
| Channel | Reaction Time | CV% | Sample Value | Channel | Reaction Time | CV% | Spike Value | Spike Recovery % |
| 1 | 404 | 0.7% | 0.528 EU/mL | 2 | 288 | 1.5% | 0.162 EU/mL | 129% |
| 3 | 400 | | | 4 | 282 | | | |

**Supplementary Figure S19**: **Endotoxin levels in plasmid sample 2 prepared using QIAGEN Midiprep Plus kit.**

| | **Plasmid 1** *From Fig. S18* | **Plasmid 2** *From Fig. S19* | **Plasmid 3** *From Fig. S17* |
|---|---|---|---|
| Plasmid yield | 380 µg | 631 µg | 180 µg |
| Sample 1 | 1.45 EU/µg | 0.628 EU/µg | 0.133 EU/µg |
| Sample 2 | 1.58 EU/µg | 0.550 EU/µg | 0.151 EU/µg |
| Sample 3 | 1.36 EU/µg | 0.528 EU/µg | 0.227 EU/µg |
| Mean | **1.46** EU/µg | **0.569** EU/µg | **0.170** EU/µg |

**Supplementary Figure S20**: **Endotoxin levels in plasmids produced using QIAGEN Midiprep Plus.** The Midiprep Plus specifies an endotoxin level below 1 EU/$\mu$g; however, this may be the best-case scenario based on the typical yield from 45 ml of bacterial culture and other growth conditions. Plasmids 1 and 2 were *p-att-ef1a-ABE8e-dCas9-BSD* (9533 bp), cultured simultaneously in identical volumes and conditions, yet differing in yield and endotoxin level. Plasmid 1 was used for the Plasmid2MC method endotoxin level reduction experiments, as it represented a plasmid with the higher endotoxin values of the two. Plasmid 2 was used for the base editing transfections. Plasmid 3 was *p-SBkit-ABE8e-parental-plasmid* (11071 bp) with the MC-Easy™ bacterial backbone, cultured with Kanamycin selection as part of the initial cloning of this plasmid.

#### Recombination efficiency as a function of ΦC31 molecules per kbp of plasmid DNA

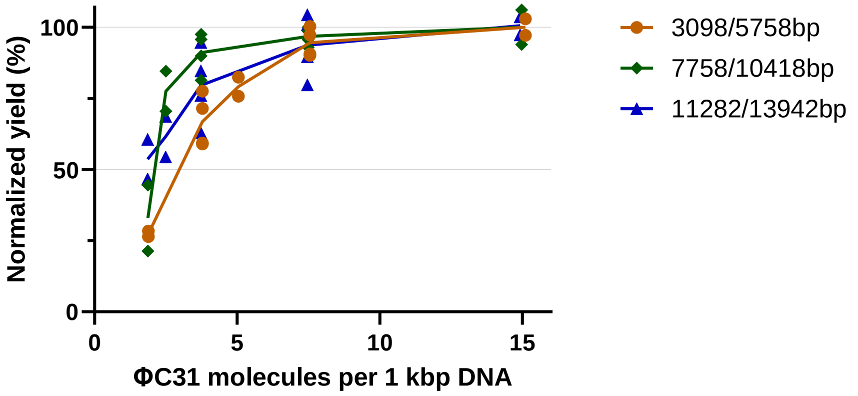


**Supplementary Figure S21**: **Recombination efficiency as a function of ΦC31 molecules per kbp of plasmid DNA.** This figure reanalyzes data from Figure 2I of the main article, converting the molar ratio to number of ΦC31 molecules per kilobase pair (kbp) of the length of source plasmid DNA. In addition to the canonical ΦC31 attP and attB sequences (which do not precisely exist in the human genome), there are approximately 27,924 pseudo-sites within the human genome that can mimic these sequences [8]. These pseudo-sites allow for the formation of functional ΦC31 dimers approximately every 107 kbp, a scenario likely extendable to any arbitrary DNA sequence. The number of sites where a single ΦC31 molecule can bind, whether transiently or more permanently, is notably higher, depending on the binding energy of each site [9]. Such off-target binding could reduce the availability of the precise four ΦC31 molecules needed to initiate recombination — two at attP and two at attB. These dynamics might account for the need for a higher molar ratio (or equivalently, a greater number of ΦC31 molecules per kbp of DNA) to achieve efficient recombination, contrasting with the theoretical ideal where just four ΦC31 molecules, specifically bound, would suffice [9].

The data points have been recalibrated for clarity so that the maximum recombination efficiencies observed in Figure 2I are normalized to 100%, with lower molar ratio data adjusted proportionately. Given that various DNA sequences might harbor multiple binding sites with different binding energies and thus different ΦC31 binding durations, the 10,418 bp plasmid achieves efficiency plateau first, followed by the 13,942 bp plasmid, and then the shortest at 5,757 bp. However, it is evident that all three plasmids begin approaching their plateau of recombination efficiency over 75% when there are more than 5 ΦC31 molecules per kbp of plasmid length, while all test plasmids exhibit diminished recombination performance at or below 2.5 ΦC31 molecules per kbp.
